# Structure-informed theoretical modeling defines principles governing avidity in bivalent protein interactions

**DOI:** 10.1101/2025.05.13.653723

**Authors:** Reagan Portelance, Anqi Wu, Alekhya Kandoor, Kristen M. Naegle

## Abstract

In signaling cascades, where domain-motif interactions tend to interact with relatively low affinity (allowing for reversibility), signaling proteins often encode multiple domains or motifs. This presents the possibility for avidity where multivalent binding drastically increases the interaction strength and duration. However, given the large combinatorial space, predicting and validating multivalent interactions that interact with avidity is a challenge. Here, we integrate mechanistic modeling, structure-based analysis, and experimental approaches as a framework for defining the conditions under which avidity plays a role. We explore the tandem SH2 domain family of interactions with bisphosphorylated partners as a multivalent archetype, which encompasses key secondary messengers in tyrosine kinase signaling networks. While certain multivalent interactions have been shown to be necessary in immune receptor recruitment of partners, bivalent recruitment of tandem SH2 domains more broadly is poorly understood. Theoretical modeling suggests that maximum avidity occurs with closely spaced tyrosine phosphorylation sites combined with moderate monovalent affinities – exactly around the innate range of SH2 domain affinity – or with phosphorylation sites separated by sufficiently flexible linkers. Surprisingly, despite sequence diversity, structure-based analysis showed relatively conserved three-dimensional spacing between SH2 domains across all tandem SH2 families, which we corroborate experimentally, suggesting evolutionary optimization for avidity interactions. The combination of structure-based analysis of domain spacing with available monovalent experimental data appears, along with iterative experimental refinement of biophysical parameters, can identify high affinity interactions of tandem SH2 domain recruitment to the EGFR C-terminal tail. Using these principles, we extended bivalent predictions into the full phosphoproteome space and structural parameterization of other partners of SH2 domain binding, providing resources and methods for more rapid expansion of bivalent analysis. These approaches lay the groundwork for larger utility in multivalent prediction and testing to help better understand protein interactions that drive cell signaling.

## Introduction

Avidity is the property that defines the increase in effective binding strength observed in interactions that involve multiple monovalent components. This occurs due to biophysical constraints where the initial interaction locks the unbound interface into a region of space constrained by their shared linker, thus increasing its “effective concentration” and, as a result, its binding driving force (1). It is a well-documented phenomenon in antibody binding, where multiple antibody epitopes boost affinity (2), which scientists have exploited for antibody-drug conjugates to increase therapeutic impact (3, 4). Cell signaling also relies on multivalent interactions to drastically increase sensitivity and specificity in network connections as we can see in a few well-documented cases involving tandem SH2 proteins that can engage with two phosphotyrosine (pTyr) sites simultaneously (5, 6). The necessity of bivalent recruitment is well documented in immune receptor-based signaling, specifically through the patterning of two pTyr sequences, such as ZAP70 recruitment to CD3*ζ* chain of the T-cell receptor (7, 8), SYK recruitment to Fc receptors for immunoglobin E (Fc*ϵ*RI), and PTPN6/PTPN11 (SHP1/SHP2) recruitment to the programmed cell death protein 1 (PD-1) and B- and T-lymphocyte attenuator (BTLA) (9, 10). In these cases, it has been shown that protein-protein interactions are high affinity, thanks to the bivalent recruitment of the tandem SH2 domain proteins and the biophysical constraints of the linkers that guide recruitment (11).

In addition to tyrosine phosphatases PTPN6/PTPN11 and tyrosine kinases ZAP70/SYK, three additional families (PI3K, PLC*γ*, and RASA1), covering ten total human proteins, also contain tandem SH2 domains. These are key proteins that transition from receptor-proximal phosphotyrosine signaling to downstream lipid signaling (PI3K and PLC*γ*) and serine/threonine signaling (RASA1). Multisite phosphorylation is also more broadly observed on other receptor tyrosine kinases (RTKs) and on downstream targets such as GAB1, which gets robustly phosphorylated to act as a docking protein for multiple tandem SH2 proteins such as PTPN11, PI3K, and RASA1, allowing for simultaneous regulation of different signaling networks (12). Beyond tandem SH2 domain containing proteins, approximately half of the SH2 domain family of proteins contain two protein interaction modules (e.g. SH3-SH2 and PTB-SH2) (13) and multidomain protein interaction modules are found broadly across the proteome. Clearly, there is strong potential for multivalency in key signaling interactions across a wide range of signaling pathways, and we are likely missing critical information by ignoring the impact of avidity on these systems. However, we currently lack the ability to predict when and how avidity will shape multivalent interactions.

One challenge to broadly identifying multivalent avidity interactions across the proteome is the complication of modeling multivalent interactions in a framework that captures the biophysics of the constrained interaction and the many species that are formed (such as an N-terminal SH2 domain bound to both pTyr sites and bivalent species bound in different orientations). Errington et al. developed an ordinary differential equation (ODE) model that accounts for the biophysical parameters of the linker constraint, using this to scale the driving force of the second interaction in the complex once the first has formed (14), which we adapted here for the purpose of exploring the theoretical regimes that define asymmetric binding.

An additional challenge in the field of tandem SH2 domain interactions is the experimental validation of a theoretical model, especially for interactions where the bisphosphorylated partners might be far apart and separated by a highly flexible linker – such as in receptor tails (e.g. EGFR). Peptide synthesis of phosphorylated proteins becomes both costly and complex, with limitations on total size and number of pTyr residues that can be incorporated (15). We developed a synthetic toolkit that can produce large amounts of multiply-phosphorylated protein, including sites across the entirety of the EGFR C-terminal tail in a single protein construct (16). Using both advances, computational and experimental, we set out to test the theoretical modeling results. Using structure-based analysis, we also explored how to inform the linker biophysics for more accurate models, finding that, despite high sequence length variability, tandem SH2 domains fall into two groups of different linker distances and despite the differences, they both can drive interactions with high avidity to similar pTyr partners. We then used an iterative approach to predict high affinity PLC*γ* bivalent recruitment across the EGFR Ctail pTyr sites, selected a large set for experimental testing, and then used measurements to better infer the linker biophysics of the EGFR Ctail and contextualize the model on its ability to separate between interactions that occur with avidity and those that do not. Based on the findings, we then sought to generalize high affinity bivalent interactions identification, involving pairs of pTyr sites and human tandem SH2 domain proteins, using available quantitative and high-throughput data, which recovers known and new protein interactions. Given the importance of parameterizing biophysical properties of linkers and domain sizes, we also undertook a comprehensive evaluation of other partners involved in multivalency with SH2 domains in tyrosine kinase signaling, identifying other generalized conservation that may guide high affinity interactions that determine signaling. Altogether, the framework presented here, and the findings in the tandem SH2 domain family, offer an approach for tackling the high complexity of identifying and testing protein interactions with emergent properties that significantly differ from their monovalent parts.

## Results

First, a computational model of tandem SH2 binding was established by modifying and expanding a previously developed multivalent receptor-ligand ODE model (14). Accuracy of this model was tested by estimating the bivalent binding affinity and avidity effect present in the PTPN11:GAB1_*Y* 627*/Y* 659_ interaction followed by experimental validation using biolayer interferometry. Once the model was validated, high-throughput simulations were performed to identify ideal parameters to optimize avidity and explore the role of avidity in increasing binding duration, and the differences across tandem SH2 protein behavior was analyzed. Moving into a more complex system, the model was used to successfully predict preferred binding partners in the PLC*γ*1:EGFR interaction, confirmed through BLI validation. Finally, the knowledge we gained in this study was applied to develop a resource that contains subsets of pTyr pairs across the human proteome that are predicted to experience avidity for various tandem SH2 proteins.

### Establishing a computational model of tandem SH2 binding

Errington et al.’s ordinary differential equation (ODE) model simulates a surface plasmon resonance (SPR) experiment between a multivalent receptor-ligand pair, tracking all possible binding configurations over time – 15 of which exist in a bivalent system (14) (Fig. S1). Key parameters in the model include the monovalent k_on_ and k_off_ rate constants, species concentrations, and linker parameters. Linker parameters include the sequence length and protein flexibility which are used to calculate the “effective concentration” of an unbound species following initial binding of the first domain, thus altering the driving force of the subsequent interaction and producing avidity. Linker flexibility is indicated by a persistence length (l_p_), where low persistence lengths suggest a more flexible protein. We adapted the model to allow monovalent affinity parameterization of all possible SH2-pTyr interactions and to improve control over the persistence length parameter. We modeled dynamic behavior under different SH2 domain concentrations, extracting the fraction bound to both SH2 domains (in either configuration) at equilibrium to estimate the bivalent dissociation constant (K_D,B_). We additionally simulated monovalent domain interactions with an SH2 ligand (by setting the other domain affinity to 1M – effectively removing that interaction). We refer to these as K_D,N_ and K_D,C_ and calculate avidity as the inverse of the bivalent dissociation constant (K_A,B_) divided by the sum of the inverse of the monovalent affinities (K_A,N_ and K_A,C_) (Equation 1):

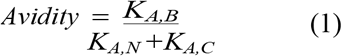

### Simulations recapitulate avidity in PTPN11:GAB1 interaction

To test the suitability of this modeling approach for tandem SH2 domain binding applications, we selected an interesting cytosolic interaction that is known to depend on avidity – PTPN11 binding to Y627/Y659 bisphosphorylated GAB1 (17, 18). The linker separating the PTPN11 SH2 domains is 9 amino acids long, and the GAB1 pTyr sites are separated by 31 amino acids. The model’s base function estimates linker length by taking the number of amino acids that encompass the linker and assuming a conservative per-residue contour increment for an extended backbone of 3Å (14). Since PDB structures exist that span the linker of PTPN11 and GAB1 (19), the biophysical separation was measured, and the linker length parameter was set to was adjusted to 5.75 a.a. and 12.95 a.a. for PTPN11 and GAB1, respectively, so that the resulting model-based separation accurately reflects the separation of 17.3Å and 38.85Å. The persistence length of the SH2 linker was set to 30Å as has been measured in interdomain spanning protein linkers (20). The pTyr linker was set to 4Å to match previously measured persistence lengths of intrinsically disordered proteins (21, 22) (Fig. 1A). Estimates of monovalent K_D_ values were taken from a deep reanalysis of fluorescence polarization experiments (23, 24) by Ronan et al. (25). The N-SH2 domain of PTPN11 is estimated to bind to site Y627 with a monovalent K_D_ of 287 nM. Unfortunately, in that specific study, data for the PTPN11 C-SH2 domain is not usable (25). We know that the C-SH2 domain is capable of binding to Y659 (17), so we modeled a monovalent K_D_ of 20 *µ*M, which is the minimal detectable binding affinity based on the experimental limits of the studies (23, 24). For simplicity, we assumed no binding of the C-SH2 domain to site Y627, to prevent interference with the N-terminal interaction. Rate constants were set with a constant k_off_ of 1 s^−1^ – approximating data found in PTPN11 and PIK3R1 SH2 binding studies (26, 27) – adjusting k_on_ for the desired K_D_. Although specific rates would be important to dynamic models, here we are specifically interested in using equilibrium behavior to understand overall binding affinity between a tandem SH2 domain and a bisphosphorylated partner, so holding k_off_ constant ensures consistency and provides further confidence in comparisons made across proteins.

**Fig. 1.**
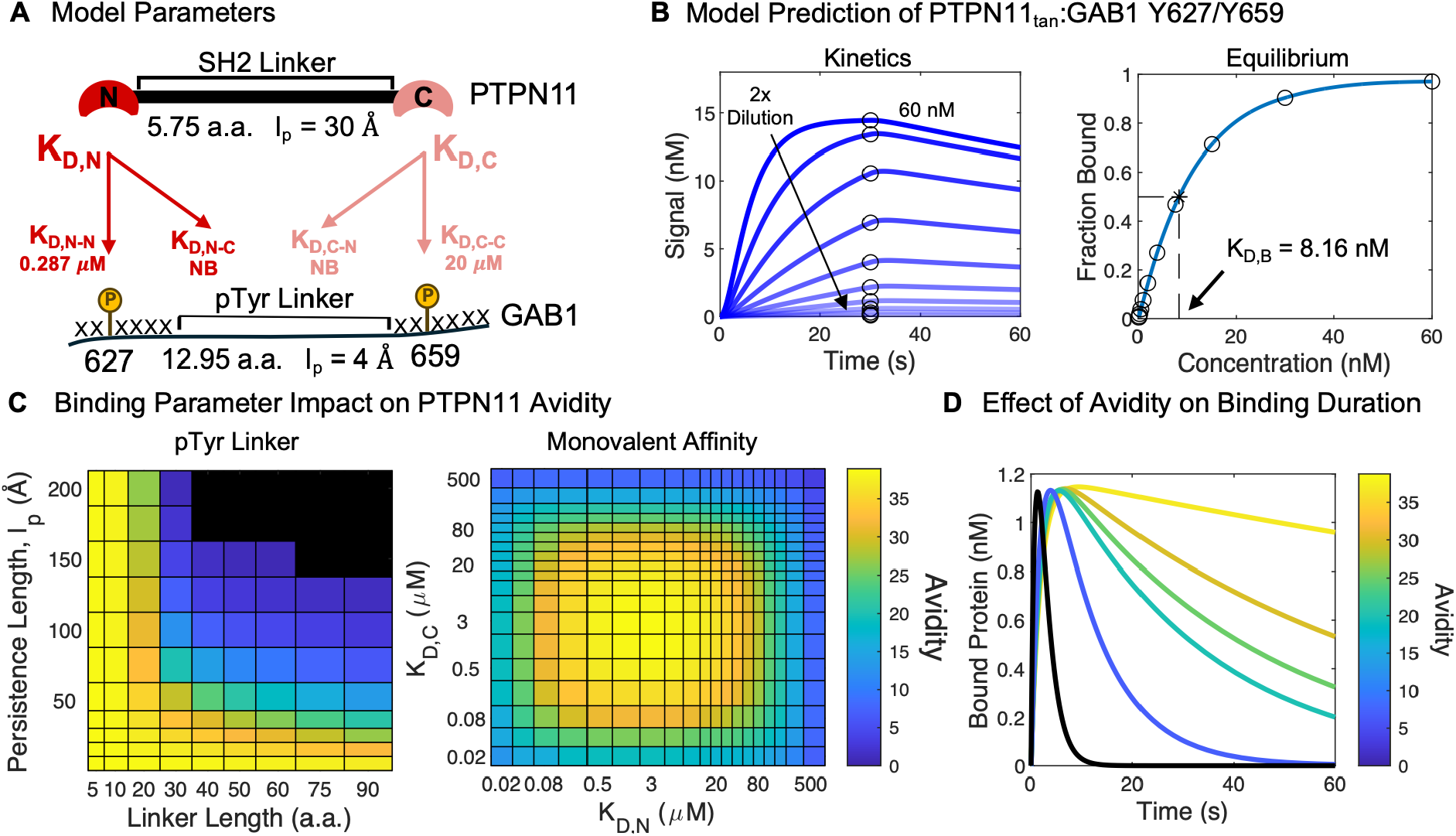
Computational simulations to determine theoretical effects of interaction parameters on avidity. **A)** The generalized ODE model considers linker parameters for both partners and the monovalent affinities of all pairs. An interaction that is not expected to occur is listed as a non-binder (NB). This toy model is annotated with parameters specifically used to model tandem SH2 domain binding of PTPN11 to doubly phosphorylated Y627/Y659 of GAB1, based on literature and sequence values. **B)** A serial dilution simulation of surface plasmon resonance (SPR) experiments of PTPN11_tan_ and GAB1 was performed using the ODE model. Equilibrium concentrations of bound protein were used to generate a concentration effect curve and estimate the K_D,B_. **C)** We used the model to independently explore the impact of ligand linker biophysics and affinity of the monovalent species. Here, the PTPN11 tandem SH2 domain parameters were used, and we either fixed monovalent affinities with changing linker parameters, or linker parameters were fixed with changing monovalent affinities. The heat map color indicates the avidity measured under each condition with black indicating an inability to achieve bivalent binding. **D)** The kinetics of transient binding (interaction does not reach equilibrium) of the SH2 domain are plotted here under different model parameters that yield varying levels of avidity. The black curve showcases the behavior of a monovalent species that experiences no avidity.

Simulations were performed with a series of eleven PTPN11 concentrations, and the resulting concentration effect curve provided a predicted K_D,B_ of 8.16 nM and avidity of 35, meaning the presence of both SH2 domains makes the binding affinity of the interaction 35-fold stronger than the individual monomers (Fig. 1B). In short, the capacity for tandem SH2 domain interaction with GAB1 converts a moderate affinity interaction (at best 287 nM) to a strong interaction of 8.16 nM, which has significant implications for the degree and duration of the PTPN11:GAB1 interaction and suggests that the modeling approach informed by available monovalent interaction data successfully recapitulates a known bivalent interaction property.

### The role of linkers on shaping avidity

Having demonstrated our ability to correctly model avidity within the PTPN11-GAB1 interaction, we wanted to explore the theoretical impact of model parameters. We first explored linker effects, since this parameter drives the biophysical constraints of the interaction. Keeping the model linker parameters for PTPN11 and fixing K_D,N_ and K_D,C_ at 4 *µ*M, we simulated a series of receptor linker and persistence lengths, while calculating avidity under each condition (Fig. 1C). We observed maximum avidity occurs in conditions where the pTyr linker is relatively short or when it is highly flexible. As the distance between the pTyr sites increases and the linker becomes more rigid, avidity decreases until bivalent binding is not possible. Across the theoretical range of linkers, it turns out that the maximum avidity attainable is closely approximated by that of the PTPN11:GAB1 interaction, suggesting that partnership is approximately optimized. It should be noted that the model cannot identify occlusion effects – cases when SH2 domains engaged with two closely spaced pTyr physically overlap. To account for this effect, we set a lower limit on the pTyr separation modeled. For the case of SH2 engagement, we hypothesized this minimum spacing is close to 12 amino acids separating the two pTyr sites (a 6 amino acid linker; Fig 1A), based on the diameter of the SH2 domain as measured across crystal structures, but in reality, it would be slightly smaller due to the SH2 domains not being perfect spheres. In support of this, the smallest bisphosphorylated ligands where we could find evidence of tandem SH2 engagement are the 10 amino acid ITAM sequences in the TCR CD3 chains which engage with ZAP70 through avidity (7, 8). Therefore, we did not model any linker below a separation of 10 amino acids, which corresponds with a linker length of 4Å.

### The role of monovalent affinities on shaping avidity

The other key parameters that determine the driving force of a tandem SH2 interaction is the monovalent affinities of each SH2 domain. To test the impact of monovalent K_D_ on avidity we fixed the linker parameters, using PTPN11 and GAB1 values, and then independently tested monovalent affinities (1C). The strongest avidities occurred when each monovalent affinity was moderately high – as either K_D,N_ or K_D,C_ became too large or too small, avidity decreased. The behavior in these two regimes is quite different, however, from a protein-protein interaction standpoint. Under high monovalent affinities, interactions are still high, and there is diminishing benefit or need for a second recruitment site. On the opposite end of the avidity drop-off, under low enough monovalent affinities, the partners are not interacting and there is no initial binding event that seeds the biophysical constraint that leads to a secondary binding event. Theoretically, the range where avidity is strongest occurs in the typical range of human SH2 domain binding (0.1 to 10 *µ*M) (28).

The primary model of avidity is each domain binding with near exclusivity to one cognate partner on the other protein with little to no cross-reactivity, such as in the PTPN11-GAB1 model. However, some bivalent interactions experience cross-reactivity where one or both SH2 domains have an affinity for both pTyr sites. To explore how cross-reactivity between partners impacts avidity, the overall K_D,N_ and K_D,C_ of an interaction were held constant at either 2.5 *µ*M or 25 *µ*M. The individual binding affinities for each SH2 domain were altered, decreasing the “direct affinity” (N-N and C-C) and increasing the “cross affinity” (N-C and C-N) to compensate and maintain the constant monovalent K_D_ (Fig. S2). Results indicate that any amount of cross-reactivity decreases the maximum avidity of the interaction. This decrease was moderate for the condition where both SH2 domains have equivalent affinities of 2.5 *µ*M with a roughly 45% drop in avidity, but the effect was more severe when one domain has a strong affinity (2.5 *µ*M) and one domain has a weaker affinity (25 *µ*M) with a nearly 70% reduction in avidity. Hence, the characteristics of high avidity include strong preferences for separate domain specificity with minimal cross-reactivity.

### The effects of avidity on binding duration

Signaling proteins bind in a transient manner, meaning they bind, unbind, and typically re-bind several times before fully dissociating and moving away within the cell (29). Since we have established that avidity within tandem SH2 proteins likely occurs due to an increased effective concentration of unbound domain upon initial binding of the first SH2 domain, it stands to reason that a multivalent protein in the process of unbinding from a target where one domain is still bound would experience the same increased driving force. In this manner, we hypothesize that protein interactions that experience stronger avidity will re-bind with a stronger affinity, thus extending the amount of time two proteins are engaged. To test this hypothesis, the PTPN11:GAB1 interaction was used as a baseline, and linker and monovalent K_D_ parameters were altered to achieve different avidities, including a simulated monovalent condition where no avidity occurs. It is worth highlighting that our selection of a constrained k_off_ of 1 s^−1^ determines the specific time scale of this simulation and the x-axis is an arbitrary unit connected to this constant. The different conditions were then used to simulate a transient binding interaction where there is no longer a constant influx of PTPN11 to maintain a steady concentration and drive the system to equilibrium, and instead the proteins are allowed to bind and fully unbind from one another (Fig. 1D). As hypothesized, the conditions that represented a larger avidity also resulted in the proteins remaining bound for a significantly longer period.

### Establishing an experimental model of tandem SH2 domain binding

Although the adapted ODE modeling approach appears to describe a known bivalent interaction well, experimental validation of model outputs is necessary, especially since there may often be uncertainty in the parameterization of monovalent affinities and linker biophysics. Therefore, we set out to establish a complementary experimental validation. We selected biolayer interferometry (BLI) for measuring binding kinetics between an SH2 domain and a phosphorylated peptide partner immobilized to a streptavidin probe. However, synthesizing multiply phosphorylated peptides is challenging and expensive, especially at the length necessary to cover physiologically relevant partners. This limitation was overcome by utilizing our lab’s SISA-KiT system which involves co-expressing a protein fused to a p40 polyproline sequence with a constitutively active tyrosine kinase fused to an ABL SH3 domain (Fig. 2A). The affinity between the ABL SH3 domain and the p40 sequence localizes the kinase to our protein of interest, driving phosphorylation and helping us achieve multisite phosphorylation. This approach has the benefit of cheaply and easily producing large batches of phosphorylated proteins, although it produces a mixture of peptide species, including nonphosphorylated, singly phosphorylated, and doubly phosphorylated partners. We set out to establish the relevancy of this experimental system and to test model predictions of tandem SH2 domain binding.

**Fig. 2.**
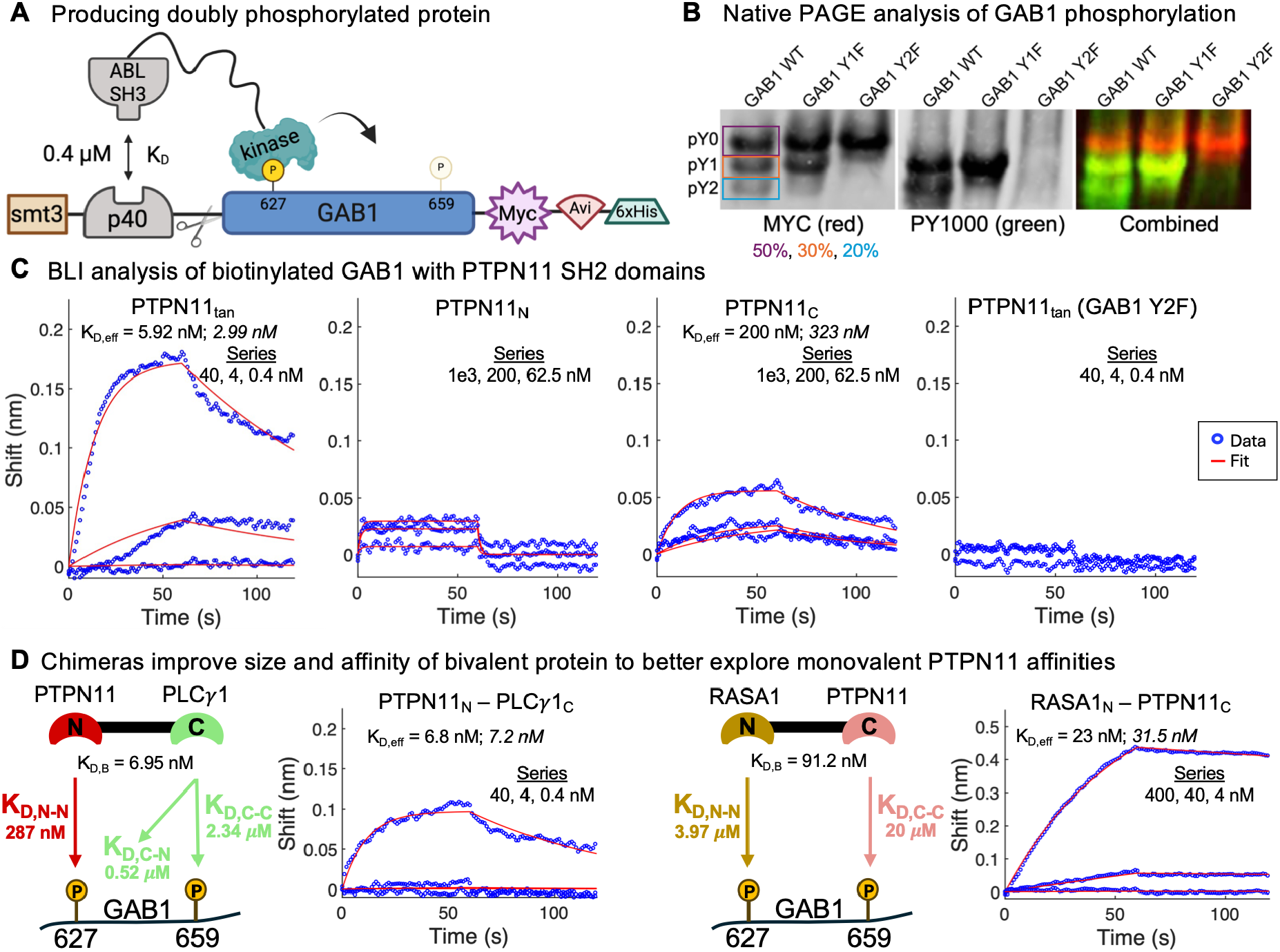
Experimental testing of the PTPN11:GAB1 interaction. **A)** SISA-KiT was used to phosphorylate both pTyr sites within the GAB1 region expressed using SRC kinase catalytic domain. The secondary interaction used in SISA-KiT to enhance phosphorylation of the substrate is the ABL SH3 domain with the p40 polyproline sequence (APTYSPPPPP). **B)** The degree of phosphorylation on purified GAB1 was evaluated by running native PAGE, immunoblotting with MYC, and confirming by pan-specific phosphoantibody. GAB1 WT refers to an unmutated GAB1 region, Y1F is a Y627F mutant (used as a reference to confirm Native PAGE behavior of singly phosphorylated species), and Y2F is a Y627F/Y659F negative control. **C)** Purified GAB1 proteins co-expressed with kinase were biotinylated using BirA ligase and streptavidin BLI tips were used to measure the binding kinetics with SH2 domain. Shown here are the background corrected on- and off-kinetics and the fit of the BLI data with a 1:1 kinetic model. We adjusted the serial dilution series to match the predicted affinity, and the specific concentrations of SH2 domain in the experiment are indicated. Where fit was sufficiently reliable in the experiment, we report an effective affinity (K_D,eff_), providing the duplicate value in italics (curves provided in Supplementary Data). **D)** Chimeras were produced by swapping out the PTPN11_N_ and PTPN11_C_ SH2 domains with RASA1_N_ and PLC*γ*1_C_, respectively. Simulations of these chimeras interacting with GAB1 were performed in the model, and BLI experiments were performed to calculate K_D,eff_ for each new species.

### Experimentally testing PTPN11:GAB1 interaction

To evaluate the general experimental approach of recombinant production of bisphosphorylated proteins using SiSA-KiT, we first evaluated GAB1 recruitment of PTPN11. We expressed the approximately 100 amino acid long span (residues 590 to 694) of the GAB1 C-terminus, where Y627/Y659 were the only tyrosines, along with a non-phosphorylatable control (Y2F) and an additional form with only Y659 (Y1F) and phosphorylated them using the SRC kinase. We used native PAGE to resolve the phosphorylated forms, estimating 20% of our GAB1 protein was doubly phosphorylated and 30% of the protein was singly phosphorylated (Fig. 2B). Prior studies in both receptor tail clustering and corollary systems (PH domains interacting with phospholipids) have suggested that presenting the active species (e.g. the bisphosphorylated species) in a background of lower to no activity species (e.g. the singly and non-phosphorylated species), helps avoid non-physiological issues such as pseudo avidity effects and issues with both over- and under-estimation of binding affinities and specificity (30–32). Hence, we chose not to perform isolation of the bisphosphorylated species, and instead tested the ability of this mixed population to reproduce relevant binding. For BLI experiments, we tested three serial dilutions of SH2 domain protein, using the Y2F control to evaluate interactions not driven by phoshorylation, and performed a global fit to measure binding affinity (ligands were biotinylated using a C-terminal AviTag™ sequence). The range of serial dilutions was adjusted to better match the affinity of each interaction, since binding saturation leads to nonlinearities and the inability to accurately extract on and off rates (Fig. S4). Like other experimental kinetic or thermodynamic analyses, BLI signal represents the mixture of all configurations of the SH2 domain protein bound to the ligand and hence we refer to the measured affinity from BLI as an “effective” affinity (K_D,eff_).

We hypothesized that if the degree of doubly phosphorylated protein was sufficient to induce bivalent binding, then the tandem SH2 domain binding would be significantly stronger than the monovalent SH2 domain binding. Hence, for this experimental test we also tested the monovalent interactions of the N-terminal and C-terminal PTPN11 domains with the phosphorylated GAB1, allowing us to calculate the avidity effect of the interaction (Fig. 2C, S3). By presenting the same phospholigand species mixture to individual N- or C-terminal domains, the measurement reflects the combined K_D,N_ and K_D,C_ interaction with both phosphorylation sites and can be compared directly to the tandem SH2 domain interaction. We performed all measurements in duplicate and provided full traces in supplements, and the K_D,eff_ values are provided for both duplicates. Here, in the text, for simplicity we will reference the interaction affinity based on the lowest K_D,eff_ observed across replicates, versus averaging values as argued in prior work (25).

Consistent with the literature values, individual PTPN11 domains bound weakly to GAB1, whereas the tandem PTPN11 domains bound strongly with an estimated K_D,eff_ of 5.92 nM, 34-fold stronger than the highest monovalent affinity, almost exactly matching the avidity predicted by the model (Fig. 2C). The bivalent affinity also closely matched values found using ITC experiments, lending additional confidence to this BLI approach (18). PTPN11_C_ bound fairly well with a K_D,eff_ of 200 nM, greatly exceeding our parameterization of 20 *µ*M. PTPN11_N_ exhibited detectable binding, but the combination of low signal and rapid on-rate made the global fit unreliable, resulting in K_D,eff_ estimates spanning six orders of magnitude for replicates with similar binding curves. Therefore, we were unable to report a K_D,eff_ for this domain. However, the bivalent affinity demonstrates strongly that both domains are functionally intact in the tandem domain species, and the prediction of the bivalent interaction was 8.16 nM, which is very similar to the experimentally measured 5.92 nM interaction. Together, these suggest that, as a set of interactions, the model affinities and the linker parameters were well calibrated to predict the overall increase in binding affinity in tandem interactions, which is two orders of magnitude better than the best monovalent affinity. Importantly, these results demonstrate that the experimental approach of producing large phosphorylated species, coupled with BLI, can measure high avidity interactions and evaluate model predictions.

### SH2 chimeras to interrogate model parameters

To overcome the signal limitations of testing low affinity SH2 monomers with BLI, we developed a new approach for evaluating the discrepancy in monovalent affinities of each PTPN11 domain. We selected two alternate tandem SH2 domain proteins, RASA1 and PLC*γ*1, which also had previously characterized binding affinities with GAB1 Y627 and Y659 (25). We made two domain “swaps” that would allow us to isolate and interrogate each monovalent PTPN11 domain under bivalent conditions that are better for BLI – increasing protein size and boosting affinity within the optimum range of the instrument (Fig. 2D). The PTPN11_N_ - PLC*γ*1_C_ chimera replaces the PTPN11 C-SH2 domain that was lacking binding affinity estimates in the literature with a new C-terminal domain with previously measurable affinity to both GAB1 sites. The RASA1_N_ - PTPN11_C_ chimera replaces the immeasurable PTPN11 N-terminal domain with an interaction previously characterized with GAB1 Y627 as 3.97 *µ*M. We simulated and experimentally tested these chimeras, keeping the originally proposed PTPN11 domain interaction parameters (Fig. 1A). For the PTPN11_N_ - PLC*γ*1_C_ chimera, the model and BLI experiments were in strong agreement with a K_D,B_ of 6.95 nM and a K_D,eff_ of 6.8 nM, suggesting that our parameterization of the PTPN11 N-SH2 domain is fairly accurate, and the monomer is binding with an affinity of roughly 300 nM. However, for the RASA1_N_ - PTPN11_C_ chimera, the model and experimental results differed by a wide margin. The model provided a K_D,B_ prediction of 91.2 nM whereas BLI yielded a K_D,eff_ of 23 nM. This disagreement suggests that the parameterization of the PTPN11 C-SH2 affinity of 20 *µ*M was inaccurate, and the monomer can bind with a stronger affinity as supported by the PTPN11_C_ BLI results. This is not surprising given the 20 *µ*M parameter was arbitrarily chosen due to limitations in literature data.

These results highlight the importance of having accurate experimental data to parameterize the ODE model. However, the strong agreement between the model and experimental results of the PTPN11_tan_:GAB1 interaction, despite the uncertainty surrounding PTPN11_C_ demonstrates the ability of the model to handle errors in one domain if the remaining monovalent interaction is relatively strong and can drive avidity. Furthermore, these results introduce the idea that certain pTyr sites within the human proteome that are not expected to interact with any SH2 domain-containing protein, based on available monovalent binding experiments, may actually be a critical binding site to enhance bivalent binding of a tandem SH2 domain partner, changing both amount and duration of the interaction.

### Exploring generalizability of tandem SH2 domain binding

Having established that the computational approach can successfully predict a known high affinity bivalent interaction with PTPN11-GAB1 and the ability to confirm experimentally, we set out to explore the generalizability of this approach to more broadly predict tandem SH2 domain recruitment. First, we asked if we could develop predictions for each family member of the tandem SH2 domains, which vary in the SH2 domain linker region. Secondly, we ask if we can scan a set of different pTyr pairs along a receptor tyrosine kinase C-terminal tail and identify the preferred binding partners of an SH2 domain.

### Expanding to all tandem SH2 domain family members

The ten tandem SH2 proteins within the human proteome are spread across five structurally homologous groups, each exhibiting a unique SH2 linker. PTPN11 and PLC*γ*1 contain a short, flexible SH2 linker of 9/10 amino acids in length whereas ZAP70 and PIK3R1 contain a much longer linker of 60/195 amino acids which is rich in alpha helices, making it significantly more rigid (the persistence length of alpha helices is typically between 150 – 200Å (33, 34)). Finally, RASA1’s SH2 domains are separated by an SH3 domain, providing moderate rigidity to accompany its length of 78 amino acids (35). When measuring the linker distance for these proteins using PDB structure estimates, the differences in size across proteins are noticeably smaller, but there still exists a modest variability – most notably between PTPN11 and ZAP70 which exhibit an average linker distance of 17.3 and 40.9 Å, respectively (Fig. 3A). We were curious whether the difference in SH2 linker parameters between ZAP70 and PTPN11 would also correspond to differences in preferred pTyr linker parameters, so we simulated a pTyr linker sweep experiment for ZAP70 (Fig 3B). Somewhat surprisingly, the results of this simulation match quite well to the same experiment performed for PTPN11 (Fig 1C), with the only real difference being a modestly improved avidity response to rigid pTyr linkers of a slightly greater length.

**Fig. 3.**
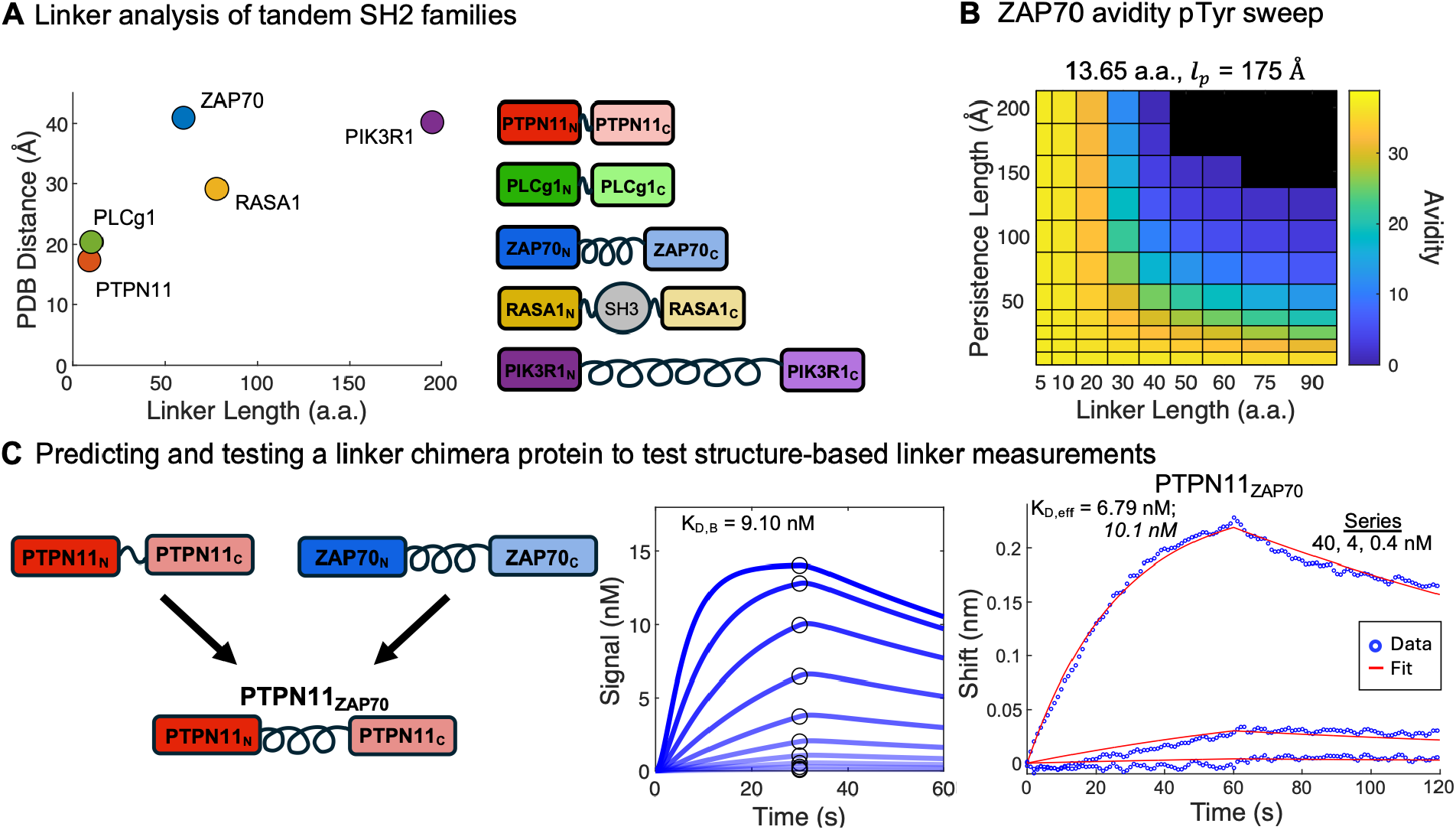
Evaluating linkers between all tandem SH2 domains across human SH2 families. **A)** The structure-based end-to-end distance versus the primary sequence length (number of amino acids between the domains) for each tandem SH2 protein family is shown. Structure-based distance is averaged across available structures for that protein. **B)** We conducted a pTyr linker sweep simulation where ZAP70 linker parameters were held constant for the SH2 linker and pTyr linker parameters were varied, with avidity being calculated under each condition. Setting the structure estimate linker length at 13.65 a.a. results in a linker distance that closely matches 40.9 Å, given the model assumes 3 Å per residue. **C)** We tested the hypothesis that structure-based distance is similar between family members, despite high sequence variability by replacing the short PTPN11 linker region between PTPN11 SH2 domains with the large, helical region of ZAP70. We modeled and experimentally measured the effective affinity of the PTPN11_ZAP70_ chimera with doubly phosphorylated GAB1 Y627/659. Experimentally measured chimera affinity (6.79nM) is very close to that of the PTPN11 binding with GAB1 (5.92nM), suggesting that the model hypothesis of similar binding characteristics across varying SH2 linker parameters is accurate.

To confirm that the variability across linkers causes minimal changes to binding characteristics of the tandem proteins, we created a linker “swap” by replacing the PTPN11 SH2 domain linker with the ZAP70 linker (Fig 3C). Based on structural parameterization, the (PTPN11_ZAP70_) chimera is predicted to bind with high avidity – on a similar scale to wild type PTPN11 (9.10 nM compared to 8.16 nM of PTPN11). Experimentally, the PTPN11_ZAP70_ bound to GAB1 with an effective affinity of 6.79 nM, closely matching the model estimate and as expected, being only slightly higher than the original PTPN11_tan_ K_D,eff_ of 5.92 nM (Fig. S5).

Given the stark differences between the flexibility of the SH2 linkers and their primary amino acid sequence lengths, it is surprising how small of a difference there is in their binding avidity with a shared target. Structurally, the longer linkers of proteins such as PIK3R1 are folded in a way that minimizes their end-to-end distance.

PIK3R1’s linker is only 2.3x longer than PTPN11’s linker, based on physical distance despite containing 22x as many amino acids, and nearly exactly matches the distance across ZAP70’s linker despite containing more than 3x the number of amino acids. This allows PIK3R1 and ZAP70 to engage just as strongly with short pTyr linkers as proteins such as PTPN11 and PLC*γ*1. These results suggest that all tandem SH2 proteins evolved a similar structural separation, which sets them up to engage with strong avidity to pTyr pairs with similar biophysical separation.

### Iterative computational and experimental assessment of PLCγ1 bivalent recruitment to EGFR

We have demonstrated the utility of the ODE model to investigate the PTPN11:GAB1 interaction, including chimeras and their impacts. Next, we wanted to test whether the framework could translate well to a new set of more complex interactions, where multiple pTyr pairs exist along an RTK with highly variable separation. We elected to study recruitment of PLC*γ*1 to pTyr pairs in the EGFR C-terminal tail, where bivalent interactions are not yet well characterized. The EGFR C-tail contains nine pTyr sites, and both SH2 domains of PLC*γ*1 have a measured monovalent affinity towards at least four of these sites (Fig. 4A). However, studies on the recruitment of PLC*γ*1 only focus on a subset of pTyr sites – completely ignoring the presence of Y998 and Y1138 – and only compare the binding affinity of individual sites (36, 37). The precise manner of bivalent engagement to EGFR is unknown, leaving a major gap in our understanding of recruitment strength and configuration which may influence the interactions between EGFR and other signaling proteins. One complication of analyzing this system is that in the past, bisphosphorylated EGFR Ctail spanning the full range has not been feasible. Complicating the modeling approach is that the flexibility and effective separation of pTyr pairs is unknown. As a starting point, we used the ODE model to predict the K_D,B_ of each potential pTyr pair and identify the rank order of preferred binding states of this multifaceted interaction, where we parameterized an initial model using a persistence length of 4Å, indicating high flexibility, and used the amino acid separation as the linker length, which as noted will be multiplied by the model’s assumed conservative per-residue contour increment for an extended backbone of 3Å (14). The two pairs spanning almost the entire length of the tail (998/1197 and 1016/1197) were not considered, since the estimation of the linker biophysics was unsolvable in our computing environment and memory. All other remaining pair predictions were ranked, based on bivalent avidity (Fig. 4B), suggesting a broad range of effective affinities, all of which indicate some degree of avidity, given this starting parameterization.

**Fig. 4.**
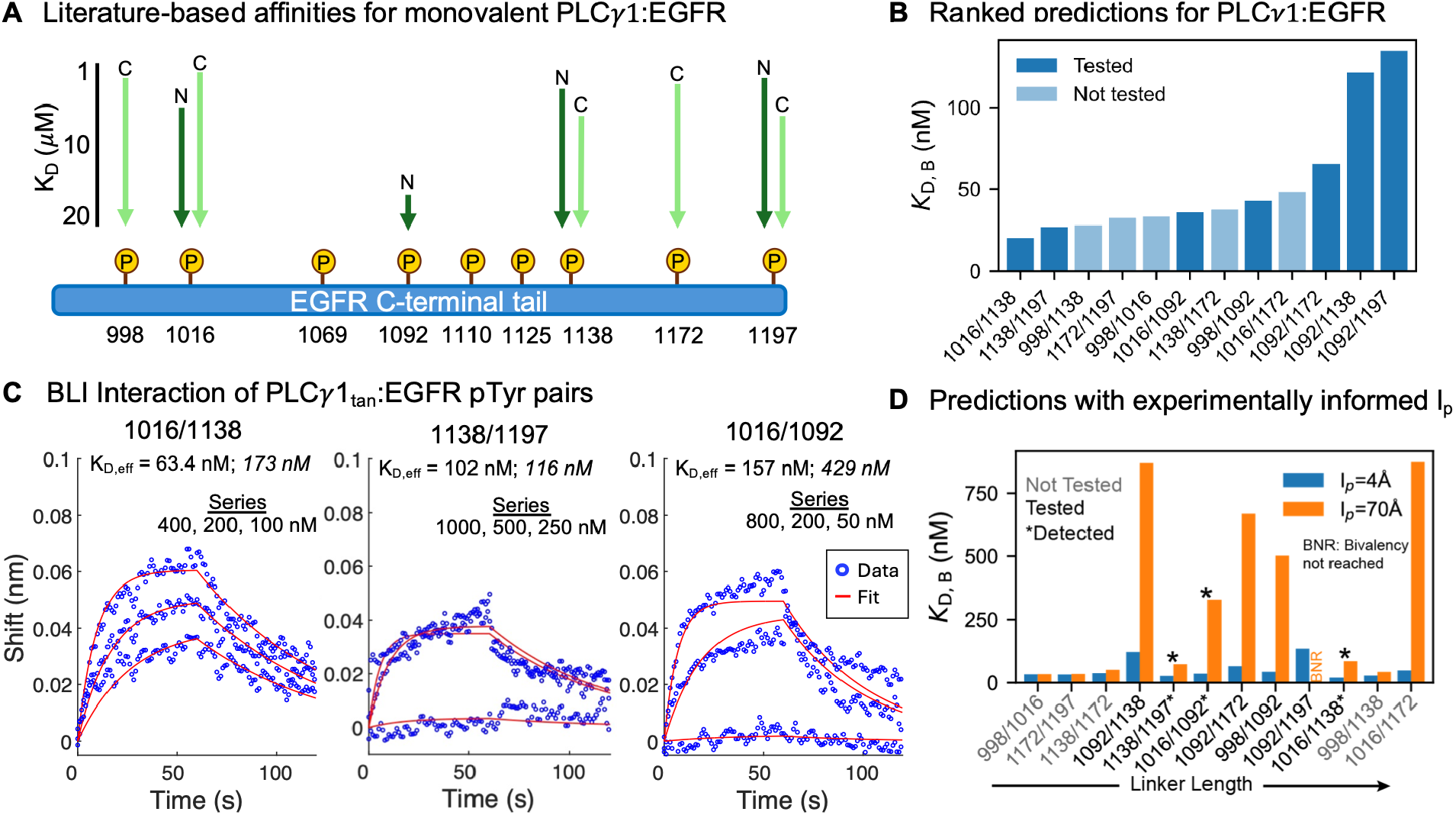
Predicting and testing bivalent recruitment of PLC*γ*1 to EGFR C-terminal tail phosphotyrosine pairs. **A)** Literature based monovalent affinities of PLC*γ*1 with pTyr sites across the EGFR C-terminal tail (25), where N is N-terminal SH2 domain affinity and C is C-terminal SH2 domain affinity, where the arrow length is proportional to affinity (no arrow indicates undetectable binding). **B)** Using literature affinity values, structure-based SH2 domain distance, a highly flexible persistence length (4A), and estimated biophysical distances for the disordered tail, we predicted the bivalent binding K_D,B_ of PLC*γ*1 to EGFR pTyr pairs (ranked by affinity). We selected seven pairs spanning the range for experimental testing. **C)** Experimental data for Y7F pTyr pairs by BLI for each interaction that had binding with avidity. All were performed in duplicate (Fig. S11) with the second replicate affinity is in italics. **D)** Original model (blue) and new model predictions using an experimentally fit lp value for the EGFRCtail, ranked by linker length between EGFR Ctail sites. The pairs tested are indicated in black text and the three interactions with avidity are starred. Increasing persistence length led to some models not reaching sufficient bivalent interaction for effective affinity estimation, indicated by BNR.

To evaluate model predictions, we selected seven pTyr pairs spanning a range of bivalent affinity predictions for experimental testing (Fig. 4B). Here, we compared Y7F EGFR C-tail constructs containing two tyrosines to a Y9F control (no tyrosines). We used a combination of Phos-Tag (Fig. S9) (Native PAGE was unresolvable for this larger protein), phosphospecific antibodies (Fig. S8), and Coomassie (Fig. S10) to evaluate the presence of phosphorylation and evaluate reagent quality. Across the experiments, we observed three pairs that yielded high affinity interactions (Y1016/Y1138, Y1138/Y1197, and Y1016/Y1092), consistent with avidity – having significantly stronger interaction than their monovalent interaction strengths measured in prior work (25). These affinities ranged from 63.4nM (1016/1138 pair) up to 429 nM (1016/1092). We also observed that replication variances were larger for the EGFR Ctail bivalent interactions (Fig. 4C), than any previous GAB1 interactions. All combinations of GAB1 experiments, including chimeras, replication was within 10nM of each other. However, EGFRCtail replications varied on the order of 100-300nM. These variances might indicate the increased complexity of the large EGFR Ctail – including ensembles of various configurations presented on the BLI tip. For the remaining four pTyr pairs we tested, these interactions all appear to occur near the limit of our system, dominated by the maximum SH2 domain concentration (1*µ*M), suggesting that these interactions do not occur with any degree of significant avidity (Fig. S12 captures many replications of attempts that show binding only at the highest concentrations of our dilution series – including altering kinases to increase sensitivity). Hence, from experiments, we concluded that three of the seven interactions occurred with avidity, producing much higher affinities than monovalent alone, and the remaining four do not demonstrate appreciable avidity, being close to the limit of our experimental system and on the same order of magnitude as the monovalent interaction values.

We noted that predictions were systematically higher than observed experimentally, both for those that demonstrated avidity and for those that interacted at affinities beyond our sensitivity limit. We hypothesized that the disparities between the model predictions and experimental findings were caused by the simplifying assumption of a high flexibility of EGFR Ctail in the initial model, given the limitation of having no structure-informed distance constraint. Hence, we next utilized the three pairs of sites that yielded measurable interactions with PLC*γ* to fit an improved linker persistence length of the EGFR C-terminal tail. Given the variable K_D,eff_ observed, we fit each individual replicate, for six total interpretations of the EGFR persistence length. The model systematically produced significantly more rigid values across all pair linkers than the initial parameter selected – ranging from approximately 50 to 100 Å. Hence, we next selected representative values within the range of a parameters estimated from experiments (60, 70, and 80 Å) and used these to re-evaluate model predictions (Table S1). Increasing rigidity systematically increases the effective affinity, which is to be expected, based on theoretical ranges explored previously. However, that change in prediction is not necessarily proportional to site separation (Fig. 4D). Although, the closest pairs of EGFR sites remain similar in the prediction of high affinity/avidity across persistence lengths (consistent with the prior theoretical exploration), the change in avidity for pTyr sites separated by greater distances does not change proportionally to that distance. Instead, the monovalent interactions become increasingly important to determining the degree of binding avidity. For example, 1016/1138, which is one of the largest separations in the tail at 122 amino acid separation, and an interaction we observed with high affinity, is predicted to be high affinity under all persistence lengths that might describe the tail (Fig. 4D and Table S1), whereas the closest pair of sites we measured (1092/1138), predicted to be very high affinity at very flexible linkers, quickly increase in predicted affinity with persistence lengths. Under experimentally-informed persistence lengths, all experiments performed are consistent with the models – the interactions experimentally observed are predicted at affinities well within our sensitivity, whereas all pairs tested that were beyond the sensitivity are consistently greater than 500nM affinities. Although the complexity of the EGFR Ctail presents challenges both computationally and experimentally, these results suggest that this general approach might be extended as a process to new complex systems, for example using an initial model to predict a high avidity interaction, measuring that interaction to improve the estimation of the regional flexibility, and then extending that to more accurately identify those interactions that are likely to occur with high avidity. Additionally, these results suggest that under pairs that are well matched to SH2 distances it may be less important to fully inform a model’s persistence length, but that highly separated pairs are capable of recruiting tandem SH2 domains with high affinity as long as the linkers are sufficiently flexible and the monovalent partnerships are amenable. Given EGFR Ctail appears to be sufficiently flexible across the length of at least 120 amino acids to recruit PLC*γ* with avidity, it seems likely that bivalent recruitment across other receptor tails may occur across large distances as well, given flexible enough linkers.

### Generalizing prediction capabilities in tyrosine kinase signaling

The combination of theoretical modeling, structure-based parameterization, and experimental approaches highlight key properties that govern high affinity tandem SH2 domain recruitment. To be more broadly extensible to predicting high affinity interactions though, the ability to identify such interactions without the need for highly detailed monovalent interaction data would be a great benefit, so we set out to compare how monovalent quantitative data and less characterized monovalent interaction data would perform for such purposes. Also, given the importance of SH2 in partnership with other domains on proteins to drive interactions, such as in partnership with SH3 domains (38, 39) or to guide tyrosine kinase targets (40), we set out to evaluate the linker and domain diameter relationships to evaluate the general conservation of biophysical properties. Here, we present new resources that identifies possible high affinity interactions and the biophysical parameterization for extending beyond tandem SH2 domain recruitment.

### Constructing a bivalent pTyr interaction resource

Using theoretical modeling and available monovalent affinities, we were able to predict and measure high affinity protein-interactions composed of bivalent pTyr recruitment of tandem SH2 domains. We next wished to explore if we could generalize these principles to identify other high affinity protein-protein interactions across the proteome as a result of tandem SH2 domain recruitment, despite limited monovalent data. For this, we first constructed a bivalent pTyr resource, guided by the principles identified from theoretical modeling – finding all pairs of pTyr sites in the human proteome that are far enough apart to allow for both SH2 domains to bind (10 amino acids) and which are likely separated by disordered segments (we excluded sites found in or separated by domains). This resource consists of 45,678 pairs of phosphotyrosine sites in the nonredundant human proteome from 3,861 proteins, meaning 18.9% of human proteins have at least one pair of pTyr sites. We observed a range of protein-level distribution of number of pairs, with 19% of proteins having between 1 and 3 pairs of pTyr, but some proteins having man (e.g. 28 proteins, including catenin delta-1 and IRS2, have more than 100 pTyr pairs). Next, we aggregated possible interactions between tandem SH2 domains and pairs of pTyr sites. Like others, we found the position specific scoring matrix (PSSM) approaches from degenerate libraries (41) failed to recover known relationships (42, 43) and so instead we used available data from immunoprecipitation (IP) experiments (44) and the fluorescence polarization data we used to parameterize prior models (25). This produces a resource that highlights possible bivalent interactions, which we then summarized to measure total number of interaction possibilities between a protein and a tandem SH2 domain – hypothesizing that, the more possible pairs that appear to match the rules of avidity, the more likely a strong protein-protein interaction exists. Known interaction partners emerge from this data, both at the individual recruitment level and the larger protein-protein interaction, suggesting this as a general approach to identify high affinity interactions. For example, GAB1 recruitment of PTPN11 to Y627/Y659 is recovered by both binding data approaches, as is GAB2 homolog recruitment of PTPN11 in the larger IP dataset. Under cumulative summation the recruitment of PTPN11 to ERBB receptors through GAB1 is evident (45), having high interaction capabilities with GAB1 and low capabilities directly with the receptors. Similarly other receptor-mediated recruitment emerges, such as preferential recruitment of PIK3R1 to ERBB3, but not EGFR (46, 47). Overall, this approach to identify biophysically possible pTyr pairs, coupled with available monovalent binding data, appears to be useful at identifying both pairwise and full protein level interactions that guide signaling outcomes and suggests the ability to more rapidly identify potential high affinity interactions driven by tandem pTyr engagement is possible without the necessity to perform highly quantitative monovalent assays.

### Biophysical analysis of other SH2 domain partnerships

Based on our findings that SH2 domains retained relatively similar physical distances, matched well to bivalent avidity across the entire family, despite higher primary structure diversity, we set out to evaluate other linkers involving SH2 domains. Using CoDIAC (13) we comprehensively extracted the domain diameters and end-to-end linker distances between SH2 domains and SH3, tyrosine kinase, and tyrosine phosphatase domains. We summarized the average values of all these partners, for use in modeling efforts (Supplementary Data S3), and visualized the relative spacing and sizes of all partners (Fig. S6). Interestingly, the size of these linkers are within a similar range as the SH2-SH2 linkers, with only the SH2-Kinase linkers of SYK/ZAP70 exceeding the maximum SH2-SH2 separation (Fig. S7), suggesting that these multivalent interactions also have the potential to be altered through avidity effects. Additionally, given domain diameters establish occlusion zones, we compared the linker distances, relative to their lengths, and the average combined diameter of a pair (Fig S7). The SH3-SH2 linkers are the smallest overall, which may suggest slightly different idealized polyproline-pTyr separation on cognate partners than what we observed for SH2-SH2 by modeling, even across highly diverse family members. Interestingly, across all tyrosine kinases we have data for, there is a conserved smaller separation between the adaptor domains, than with the kinase catalytic domain. Where SH3-SH2 physical separation is highly conserved, we observed much higher diversity when SH3 domains follow the SH2 domain, suggesting higher diversity in possible partner spacing between the pTyr and the polyproline, or possibly more frequently involved in trans-interactions. Overall, the analysis identifies interesting constraints that could be used to identify partner bivalent motifs and, like SH2-SH2 domains, several well conserved properties which could simplify the identification of partnerships.

## Discussion

A guiding principle in the field of phosphotyrosine signaling is that the relatively low affinity of SH2 domains (on the order of *µ*M) is important to the reversibility of signaling – allowing the dissociation of signaling complexes and phosphatase access (48–50). This study, along with existing knowledge in the immune receptor system, indicates that tandem SH2 domain containing proteins can be recruited with a strikingly higher affinity (on the order of nM). Both total amount and dynamics shape cellular outcomes (51, 52), and increasing this by 30- to 100-fold has profound impacts on protein interactions and downstream consequences. Notably, we found that the normal range of affinity for an SH2 domain (0.1 -10 *µ*M) matches well to the range of monovalent K_D_ values that maximize avidity, providing a new perspective on SH2 binding affinity – perhaps the relatively low affinity of SH2 domains evolved because it optimizes the advantage of multivalency within the tandem SH2 proteins to allow for these dramatic improvements in affinity. Additionally, we found it interesting that the tandem SH2 domain families appear to have evolved relatively similar geometries in 3D structure that set them up exceedingly well for high avidity interactions at these monovalent affinities. However, we also found that high avidity interactions were possible under much larger separation than those matched to the SH2 domain separation. For example, EGFR C-terminal tail sites that were 120 amino acids apart recruited bivalent PLC*γ* with high affinity, even across reasonable estimations of EGFR persistence lengths (an experimental finding that would not have been possible via peptide synthesis methods). Given the breadth of signaling proteins covered by the tandem SH2 domain family, there is a possibility that almost all phosphotyrosine signaling is governed to some degree by tandem SH2 domain interaction avidity. Additionally, if effective affinities are really in the nM range, then active forms of reversibility may be necessary and might explain findings in the regulation of SH2 domain linker regions (11) and on or near binding interface residues of the SH2 domain (13). Finally, the modeling approach also highlights the importance of particular mutation effects – where loss of the necessary seed sequence interaction might be more important and predictive of dysregulated interactions, then impacts on the lower affinity partner.

Given the high degree of multidomain interaction units throughout the proteome, it is likely that broad swaths of molecular biology research should consider avidity, yet the combinatorial complexity of these systems have created barriers to such avidity incorporation. Combining mechanistic modeling and structure informed parameterization presents a broader framework for identifying protein interactions with high effective affinities due to multivalency, and it provides theoretical constraints that would help identify those interactions that experience avidity. Even with limited information about monovalent species or details of linker biophysics, modeling helps to bound the reasonable space of whether high affinity interactions might occur and rank-order binding partner preference, even if not able to perfectly predict effective affinity. In this study, we also found that the iteration of modeling and experiments could help improve the efficiency of research. For example, in the future, given what we learned in the EGFR Ctail experiments, we would test ranked hypotheses across pairs in a protein, then use the fitted persistence length of the first successful experiment to reevaluate future pairs and experiments.

Additionally, the synthetic toolkit for driving tyrosine phosphorylation on proteins can easily be extended to cover alternate post-translational modifications and mixtures of bivalency (e.g. polyproline and a pTyr for SH3-SH2 testing and possibly acetylation recruitment of tandem bromodomains). It appears that these constraints can then be used to scan the proteome for possible cognate partners, using less quantitative approaches (such as immunoprecipitation) to identify the seed interaction that provides the biophysical driving force of high affinity interactions. Conservation of biophysical parameters across families, such as SH3-SH2 and SH2-SH2, suggests that the problem might be reduced for the entire family to the same patterning on cognate partners, simplifying the modeling and prediction processes. Of course, even though modeling and experiments of these two-part systems suggest important findings for signaling consequences, it does not yet consider competition, cellular localization constraints, or other dynamic processes, such as conformational dynamics that regulate different states of multidomain proteins, that are also at play in physiological signaling systems. Despite this, the framework presented here offer an approach for tackling the high complexity of identifying and testing protein interactions with emergent properties that significantly differ from their monovalent parts, which is an important step in modeling and dissecting complex interactions that govern cell physiology and considering more broadly how to intervene in dysregulated signaling cascades in disease.

## Materials and Methods

### Cloning and mutagenesis

#### Substrate and kinase vectors for phosphoprotein production

We used SiSA-KiT (16) (https://github.com/NaegleLab/SISA-KiT), which includes an SH3-fused tyrosine kinase and a polyproline-fused substrate vector, co-expression, and substrate purification as outlined further below to produce phosphorylated substrates. We cloned GAB1 and EGFRCtail into substrate vector sVd-Avi (a pGEX backgone with GST removed and an AviTag, Myc, and 6xHis inserted) containing an N-terminal p40 polyproline targeting sequence (APTYSPPPPP), using Prescission cleavage to remove the targeting sequences before downstream analysis. We have deposited this as part of the SiSA-KiT vector options with at Addgene Plasmid # 259390 – sVdAvi-GAB1. We used the following kinase vectors in this study: kVh-SRC (Addgene plasmid # 259387), kVh-ABL (Addgene plasmid # 259379), kVh-LYN (Addgene plasmid # 259385) and kVh-EPHB1 (Addgene plasmid # 259380). GAB1 – from pCDNA3.1-HA-GAB1 which was a gift from Dr. Matt Lazzara (53). EGFR C-terminal tail was gifted to us by Linda Pike in both Y9F and Y8F formats (54). We selected the appropriate Y8F plasmids to mutate a second tyrosine to phenylalanine to create the Y7F plasmids (mutagenesis described below). Using cloning processes outlined below, we cloned the span of GAB1(UniProt ID Q13480) from amino acid 590 through the end of the protein, which contained only two tyrosines (Y627 and Y659) and EGFR (UniProt ID P00533) C-tail from amino acid 980 through the C-terminal end.

#### Cloning, and mutagenesis

We used either restriction/ligation cloning or In-Fusion cloning for engineering of backbone vectors, insertion of tags and targeting. We used Snapgene to design primers, and sequenced verified by Sanger sequencing. Sequencing was performed to get full coverage of insert. Some constructs were whole plasmid sequenced to verify the entire plasmid. For cloning, we used NEB Phusion polymerase (part no. M0530S) and included extensions for cloning into the target backbones using NEB enzymes. We used Agilent’s QuickChange mutagenesis kit (part no 210518) to perform point mutations. We used carbenicillin at 100*µ*g per mL for selection of ampicillin plasmids (substrate and GST plasmids). Subcloning Efficiency DH5*α* Competent Cells: ThermoFisher Scientific Cat. 18265017 were used for DNA propagation. SH2 domains were cloned into a pGEX backbone. The sources for SH2 domains were: PTPN11 – from pBABE-PTPN11 which was a gift from Dr. Matt Lazzara (55); RASA1 – pDONR223_RASA1_WT was a gift from Jesse Boehm & William Hahn & David Root (Addgene plasmid # 81778 ; http://n2t.net/addgene:81778 ; RRID:Addgene_81778); ZAP70 – pDONR223-ZAP70 was a gift from William Hahn & David Root (Addgene plasmid # 23887 ; http://n2t.net/addgene:23887 ; RRID:Addgene_23887); PLC*γ*1 – pGEX PLCg1(NC)-SH2 was a gift from Bruce Mayer (Addgene plasmid # 46471; http://n2t.net/addgene:46471 ; RRID:Addgene_46471).

### Protein expression and purification

#### Protein induction and bacterial lysis

**SH2 Domains** Plasmids were transformed into BL21 gold DE3 (Agilent part no. 230132) and an isolated colony was used to inoculate a 5 mL LB, 5 *µ*L carbenicillin culture which grew overnight at 37^*°*^C. 2.5 mLs of this culture was used to inoculate a 100 mL LB, 100 *µ*L carbenicillin culture which shook at 37^*°*^C until it reached the desired OD600 (0.7-1.0). The culture was then induced using 0.5 mM IPTG and grown overnight in an 18^*°*^C shaker. The culture was spun down at 8500 rpm for 10 minutes and the pellet was frozen at - 20^*°*^C until lysis.

**Phosphorylated Proteins** Plasmids (substrate and kinase) were co-transformed into BL21 gold DE3 (Agilent part no. 230132) and an isolated colony was used to inoculate a 5 mL LB, 5*µ*L carbenicillin/kanamycin culture which grew overnight at 37^*°*^C. 2.5 mL of this culture was used to inoculate a 100 mL LB, 100*µ*L carbenicillin/kanamycin culture, which shook at 37^*°*^C until it reached the desired OD600 (0.7-1.0). The substrate protein was then induced using 0.5 mM IPTG and grown overnight in an 18^*°*^C shaker. The culture was spun down at 8500 rpm for 10 minutes and resuspended in 100 mL LB, 100 *µ*L carbenicillin/kanamycin, and 0.2% L-arabinose. The culture was grown for another 4 hrs in a 37^*°*^C shaker and then spun down at 8500 rpm for 10 minutes and the pellet was frozen at -20^*°*^C until lysis.

**Lysis** We resuspended bacterial cell pellets in a 10:1 ratio of final growth volume to lysis buffer. Lysis buffer was 50mM Tris-HCl at pH7.0 and 150mM NaCl, supplemented with protease inhibitors (EMD Millipore 539137), phosphatase inhibitors (Millipore Sigma P5726), and PMSF (1:1000). For small volume lysis of 1mL to 5mL, we used bead beating, adding 400*µ*L of beads to each 1mL of resuspended cell pellet and bead beating for 2 minutes. For larger lysis volumes, we sonicated resuspended pellets for 5 minutes with 40% pulse sequence. Following lysis, we clarified by centrifugation at 13,000 rpm for 10 minutes at 4^*°*^C.

#### Protein purification

**Nickel/6xHis Purification**. We used Genscript Ni-NTA MagBeads (cat no. L00295), using a magnetic tube stand for isolating beads during decanting. Prior to protein binding, we equilibrated beads in lysis buffer (50mM Tris-HCl, 300mM NaCl (pH 8.0), 0.5% TX-100, 80 mM imidazole). Beads were isolated by magnet and decanted. We added equilibration buffer again, this time in addition to 1mL of clarified lysate and incubated at 4^*°*^C for at least 1hr up to overnight, while rotating. Isolated beads were washed in a 20-fold excess (relative to bead volume) of wash buffer (50mM Tris-HCl, 300mM NaCl, 40mM Imidazole, 0.5% Triton-X (pH 8.0)) a total of three times, incubating in the wash buffer for 5 minutes each time. PreScission protease (GenScript cat no. Z02799) was used to cleave the N-terminal solubility tag and p40 sequence. The beads were resuspended in 50 mM Tris-HCl, 150 mM NaCl, 3 mM DTT at 4 times the resin bead volume and incubated overnight at 4^*°*^C with 1 *µ*L of PreScission. Protein was eluted using 2 times the resin bed volume with elution buffer (50mM Tris-HCl, 300mM NaCl, 500mM Imidazole pH 8.0). We repeated elution two total times. We dialyzed protein following manufacturer directions for the Pierce Slide-A-Lyzer mini dialysis with a 3.5kDa cutoff into 50mM Tris-HCl, 150mM NaCl at pH 7.8. Proteins were aliquoted and frozen at -80^*°*^C.

**GST Purification** We used Pierce Glutathione Agarose (part no. 16100) and followed the manufacturer’s base protocol for GST-based purification. Beads were equilibrated in buffer (50mM Tris, 150mM NaCl) and protein capture was performed for one hour to overnight at 4^*°*^C. Beads were washed (three times) with a 10-fold excess volume of equilibration buffer, with centrifugation to isolate the beads. We used PreScission protease-based elution according to manufacturer’s directions (GenScript GST-PreScission protease part no. Z02799), incubating protease with beads resuspended in 50 mM Tris-HCl, 150 mM NaCl, 3 mM DTT overnight at 4^*°*^C. PreScission cleavage was performed to ensure no dimerization of the GST domain (49).

#### Biotinylation

Following nickel-purification and dialysis, phosphorylated substrates were biotinylated using the BirA500 kit (Avidity EC 6.3.4.15). We reacted 10nmol of substrate with 2.5*µ*g of BirA ligase for between 30 minutes and 5 hours at 30^*°*^C depending on the protein concentration. Proteins were dialyzed following biotinylation to remove excess biotin. Biotinylation was frequently confirmed through 680IRdye-conjugated streptavidin, incubated for one hour at 4^*°*^C (at 1:1500), prior to final washes and LICOR-based imaging.

#### Western-based analysis

Samples were reduced in Laemmli loading buffer (Boston BioProducts) and boiled for 10 minutes at 95^*°*^C, unless they contained imidazole, where we boiled at 70^*°*^C for 10 minutes to avoid breaking protein bonds. We performed standard SDS-PAGE analysis using purchased precast NuPAGE gels from Invitrogen, typically using 4-12% gradients or single 10% gels, running in a MOPS SDS running buffer (Novex Invitrogen). Transfer to nitrocellulose membranes occurred in a buffer (Novex Invitrogen) with 20% methanol (membranes were pre-wetted in transfer buffer). Pre-stained ladder from LI-COR (Chameleon Duo) was used along with IRDye-680 and -800 secondary antibodies for infrared scanning. We used Intercept Blocking Buffer (LI-COR), diluted 1:1 in Tris Buffered Saline (TBS) for blocking membranes (one hour at room temperature or overnight at 4^*°*^C) and preparing primary and secondary antibodies. Following primary incubation (1 hour at room temperature or overnight at 4^*°*^C) and secondary incubation (1 hour at room temperature at 1:10,000, we washed membranes in a TBS-T (1% tween solution) or TBS solution. Antibody stripping was done with 0.2N NaOH for 30 minutes, as needed for efficient stripping. Membranes were scanned on a LI-COR Odyssey system.

#### Antibodies

We used the following antibodies for verification of expression and phosphorylation: MYC antibody Mouse Bio X Cell BE0238 at 1:3000 dilution; Rabbit PY1000 antibody CST 8954S at 1:1500 dilution; EGFR pY998 Rabbit CST 2641 Lot 3 at 1:1000 dilution; EGFR pY992/EGFR pY1016 Rabbit CST 2235 Lot 11 at 1:1000 dilution; ERBB2 pY1196 (with target detection of EGFR pY1138 as shown in Ryan et al. (16)) Rabbit CST 6942 Lot 1 at 1:1000 dilution; EGFR pY1068/EGFR pY1092 Rabbit CST 3777 Lot 17 at 1:1000 dilution; EGFR pY1148/EGFR pY1172 Rabbit CST 4404 Lot 4 at 1:1000 dilution; EGFR pY1173/EGFR pY1197 Rabbit CST 4407 Lot 9 at 1:1000 dilution. We used the following secondary antibodies from LI-COR Biosciences (at 1:10,000 dilution): Donkey

Anti-Mouse IgG 680RD, Donkey Anti-Rabbit IgG 800CW; Donkey Anti-Rabbit IgG 680RD; Donkey Anti-Mouse IgG 800CW. Phosphospecific antibodies were proven to be phosphorylation-dependent, as they showed no detection on constructs produced without kinase expression (negative controls). In other work (Ryan et al. (16)) we tested the cross-reactivity of these EGFR and ERBB2 antibodies with each of the EGFR sites that are part of our tandem pairs and the only cross-reactivity we observed was ERBB2 pY1196 detecting EGFR pY1016, in addition to EGFR 1138 and small reactivity of the EGFR pY1148 antibody with other sites, which is consistent with what we observed in this study with two site presentation of the EGFR tail.

#### Native PAGE

Phosphorylation efficiency of phosphoproteins was evaluated by running protein samples by native PAGE to allow for separation of protein species with varying levels of phosphorylation. A 9% acrylamide separating gel was poured with 1.8 mL 30% acrylamide, 1.5 mL 1.5 M Tris-HCl, 600 *µ*Ls 10% APS, and 6 *µ*L TEMED. A stacking gel was poured using 533 *µ*L 30% acrylamide, 1 mL 1.5 M Tris-HCl, 40 *µ*L 10% APS, and 8 *µ*L TEMED. Gels were cast using the Bio-Rad Mini-PROTEAN® Tetra Handcast System. 1L of 1x Running Buffer (250mM Tris-HCl and 1.92M Glycine) was used, and gels were run at 4^*°*^C at 90V until separation was achieved. Samples were prepared in loading dye: 40% Glycerol, 248 mM Tris-HCl, 0.02% Bromophenol Blue. Gels were analyzed using LICORbio Empiria Studio software. The MYC-labelled bands were isolated and their total signal quantified. The percentage of doubly, singly, and non-phosphorylated GAB1 protein was calculated by summing the MYC signal for all three bands in the gel and taking ratios of each individual band to the total.

#### Protein concentration estimation

We used PAGE-based separation and Coomassie staining (ThermoScientific Pierce Coomassie Brilliant Blue G250 part no 20279), followed by 680-infrared detection by LICOR for estimating protein concentration compared to a BSA standard curve.

#### Biolayer Interferometry (BLI)

Experiments were run on the Gator® Pilot BLI instrument using Streptavidin (SA) probes (Gator SKU 160002), MAX plates (Gator SKU 130062) for probe loading, and black flat plates (Gator SKU 130150) for sample loading. A sampling rate of 10 Hz was used, and a temperature of 30^*°*^C was maintained within the instrument. 250 *µ*L of buffer (50 mM Tris-HCl, 150 mM NaCl, 1% Bovine Serum Albumin, filter sterilized) was loaded into the MAX plate to allow for equilibration of the probes, and a 10-minute, 400 rpm equilibration step was performed at the beginning of each experiment. 180 *µ*L of sample was loaded into the black flat plate and held at a tilt throughout the experiment. Experiments were run with a 30 second baseline in buffer, 120 second load of biotinylated, phosphorylated protein, 30 second baseline in buffer, 60 second association of SH2 domain, and 60 second dissociation in buffer. The load wells contain equal concentrations of phosphorylated protein, whereas the association wells contain a serial dilution of SH2 domain within the first three wells of the column. The final well contains buffer to act as a reference. Plates were shaken throughout the experiment with PTPN11:GAB1 experiments being shaken at 1000 rpm and PLC*γ*1:EGFR experiments being shaken at 400 rpm.

#### BLI Analysis

Kinetic analysis was performed within the Gator One® software (version is 2.16.6.0130). Data was aligned on the Y-axis at the association step, and an inter-step correction was applied along with Savitzky-Golay filtering to reduce noise. The reference well is subtracted from the other three wells to account for non-binding artifacts within the signal. A 1:1 global binding model is fit to both the association and dissociation curves of each set of three serial dilutions with an unlinked Rmax to quantify the dissociation constant, K_D_. BLI figures contained within this paper uses data processed in the manner described above, and both the data and binding fit curves have been re-plotted in MATLAB®. The data is subsampled to include every 10th data point. All BLI data is provided on Figshare at 10.6084/m9.figshare.31141453, which was not subsampled but did undergo the other pre-processing steps.

#### ODE Modeling of tandem SH2 domain interactions

The original code was written by Dr. Wesley Erring-ton in the Sarkar lab at the University of Minnesota (14) and shared by compressed folder by Dr. Sarkar. The code was adapted to include greater flexibility within the parameters. Most notably, additional k_on_ and k_off_ parameters were incorporated into the differential equations to allow for different affinities by each SH2 do-main for each pTyr site, and persistence length was made universally adjustable instead of binning all protein interactions into two possible groups, flexible and rigid. The output of the original code is a series of plots mapping the binding behavior of the proteins under different initial concentrations of SH2 domain. Fifteen binding configurations are possible within a bivalent system, but for the purposes of this paper, only subsets were summed together and plotted as a single metric of binding. When analyzing tandem SH2 domains, the two bivalently bound configurations (inline and twisted) were isolated, whereas configurations detailing single domain binding were utilized when simulating monovalent binding. Additional code was written to extract the equilibrium concentration of each of these plots and feed that into a concentration effect curve plot. This operates by plotting the fraction of bound species at equilibrium – calculated by dividing the equilibrium concentration by the starting concentration of phosphorylated protein (R0) – against the starting concentration of SH2 domain (L0). Within the model, the concentration of SH2 domain is assumed to remain constant throughout the simulation. The value of L0 that corresponds to 50% of the phosphorylated protein being bound at equilibrium is the dissociation constant (K_D_). Results provided were generated using Matlab R2024a or R2024b. All parameters have been provided in Data S1. We used the University of Virginia’s High-Performance Computing environment Rivanna for model predictions. All Matlab code is provided in our repository at https://github.com/NaegleLab/BivalentModeling_tanSH2.

#### Estimation of diameters from structure

For simulations provided in this paper, estimates of the molecular weight of SH2 domains and pTyr sites were calculated using the following website https://web.expasy.org/compute_pi/. For pTyr sites, the sequence spanning two sites N-terminal of the pTyr site and four sites C-terminal was used to estimate molecular weight. Sites Y627 and Y659 within GAB1 were calculated and used as an estimate for all pTyr pairs.

#### SH2 diameters

We identified PDB entries containing the domain of interest (SH2) and their corresponding domain boundaries using CoDIAC (13). We included wildtype and mutant structures only if the mutations resided well outside the domain regions. To estimate domain diameters, we assumed each domain adopts an approximately spherical shape. Based on this assumption, the diameter was defined as the longest distance between any two atoms within the domain. To compute this, we measured all pairwise distances between backbone atoms (N and C) within the defined domain boundaries and recorded the maximum distance as the domain’s diameter for multiple PDB structures available for the same domain. Since the ODE code assumed the same size for both entities, and since we saw relatively small differences between the N and C terminal diameters, we averaged all diameters for both domains across all available structures. All diameter calculations are provided in Data S2. Code for extraction of domain diameters is provided at https://github.com/NaegleLab/BivalentModeling_tanSH2.

#### Estimation of linkers from structure

pTyr linkers Linker length was estimated as the number of amino acids between the pTyr sites or SH2 domains, unless a PDB file was available for the protein in which case linker length was estimated using the same PDB-extraction based approach as SH2 domains.

#### SH2 domain linkers

Structures exist for all tandem SH2 domain family members and so we used those to estimate the linker lengths. For SH2-SH2 linker used for model simulations within the paper, we used UniProt defined SH2 domain boundaries to define the linker (as the first amino acid after the N-terminal SH2 domain and up to the amino acid before the start of the C-terminal SH2 domain). Then for all available structures, we calculated the average distance between all atoms of the two amino acids farthest apart from each other on the linker region (i.e. the residues immediately adjacent to the domains). All linker calculations are provided in Data S2. Code for extraction of linker distances is provided at https://github.com/NaegleLab/BivalentModeling_tanSH2.

#### SH3, PTP, and Kinase linkers

For the expanded analysis of SH2-SH2, SH2-SH3, SH2-PTP, and SH2-Kinase linkers, we first identified all available structures containing domains of interest using CoDIAC (13). For each structure, domain diameters and inter-domain linker distances were computed as described previously. For this analysis, all distance measurements were calculated using the backbone reference point ‘CA’ atom. Domain and linker regions were defined using InterPro annotated domain boundaries. The primary analysis focused on adjacent domain pairs connected by linkers of defined amino acid lengths. In addition, we included the non-adjacent N-SH2 and C-SH2 domain pair in RASA1, where an SH3 domain lies between the two SH2 domains. Structures bound to inhibitors were excluded, as inhibitor binding often stabilizes inactive conformations that differ substantially from native states. Structures containing unmodeled regions within either domain or at residues used to define linker boundaries were also excluded. For each domain-domain pair, we assessed the consistency of linker distances and domain diameters across multiple structures. Structures exhibiting outlier behavior relative to the group were excluded from this analysis. For example, the SH2-Kinase linker distance in SRC structure (PDB: 1Y57) was at least 20Å longer than in the other SRC structures (Fig. S13B) and was therefore excluded. Similarly, several HCK structures (PDB: 2C0T, 2C0O, 2C0I) that displayed unusually large SH2 and SH3 domain diameters were excluded (Fig. S13A). Overall, SH2-Kinase domain-linker measurements from all HCK structures showed poor agreement with those from its close homologs SRC and FES (Fig. S13C). Consequently, all the SH2-Kinase measurements from HCK were excluded, whereas SH3-SH2 measurements from HCK that were comparable to those of SRC and FES were retained (Fig. S13C). For all the structures included in this analysis, domain diameters and linker distances are provided in Data S3. This includes both a comprehensive list of structures retrieved from CODIAC as well as a filtered list that was used in the final analysis and a summary of unique domain-domain pairs with average domain diameters and linker distances. All code and data sources are available at Figshare at 10.6084/m9.figshare.31141453.

#### Estimation of monovalent KD values

Estimates of monovalent K_D_ were taken from Ronan et al. meta-analysis of fluorescence polarization experiments (25). All k_off_ values were set to 1 s^−1^, and k_on_ values were adjusted to match the estimated K_D_.

#### Bivalent pTyr resource generation

We used ProteomeScoutAPI v3.0, with data updated as of January 2026, using the indicated non-redundant reference proteome as determined by Uniprot to identify all pairwise pTyr pairs in the unique human proteome (20,405 protein records). We excluded PTMs that occur within InterPro domains. We then built a pairwise pTyr resource, excluding any pairs that were less than 10 amino acids apart and any pairs that had a defined InterPro domain between them. Next, we took the two binding resource datasets, Martyn et a. (44) and Ronan et al. (25) and used the Uniprot ID and the indicated sites to map those into our reference phosphoproteome and indicated in the reference bivalent if an interaction was detected between a tandem SH2 domain and one of the sites in the pair. For Ronan et al. data, we mapped interactions if the monovalent K_D_ was less than 10*µ*M. We excluded SYK and ZAP70 mapping since most of those data were indicated as non-functional. Additionally, only the PTPN6-C domain was available in the experiment and half of the interactions tested also were indicated as non-functional SH2 domain and we excluded the mapping of these interactions. Given the highly quantitative and SH2-extensive nature of the FP data, we tracked and noted a partnership between a pair of sites and a tandem SH2 domain if at least one interaction was found and tracked the most likely configuration, twisted or inline, based on the highest sum of affinity values. For Martyn et al., we mapped each dataset that was pulled down by a tandem SH2 domain into our reference proteome, using the stimulated form (pervanadate treated) of the dataset from the publication. Martyn et al. performed IP with both wild type domains and superbinders – SH2 domains engineered to deepen the pocket, which has been shown to increase affinity without substantially altering specificity (13, 44, 56) – we tracked and noted whether the partnership was seen in one or the other type of IP – indicating interactions seen in wild type domain pull downs as HA for “high affinity” and in superbinder pull downs as LA for “low affinity”. Using the Martyn pulldown data, we were able to annotate 622 human proteins, or 45% of the human pTyr bivalent proteome with information about PIK3R1/R2 and PTPN6/11 interactions and with Ronan we annotated 406 pTyr pairs across the majority of the tandem SH2 domain family. The full resource, including all pTyr pairs, annotated Ronan data, annotated Martyn data, and summarized protein-protein interaction data is provided in Supplementary Data S4 and the code used to construct, annotate, and summarize along with primary data resources is provided on Figshare at 10.6084/m9.figshare.31141453.

#### OPAL scoring of bivalent pTyr reference

We attempted to use degenerate library-based scoring to annotate recruitment. Based on structural analysis from Kandoor et al. (13), which identified consistent -2 to +4 interactions, we scored sequences using just the -2 to +4 interaction matrix and we removed from the reference pTyr proteome any N- or C-terminal interactions that were shorter than the span provided and then we performed PSSM matrix multiplication and summation, as per (41), to score each peptide using matrices as published in Yaron-Barir et al. (57). We tested both a percentile cutoff and a score cutoff to indicate likely interactions on the real reference phosphoproteome and a random proteome, where we randomly sorted the 20 amino acids surrounding the central pTyr residue and rescored. We assessed individual and cumulative interactions across all possible scoring approaches and failed to see physiological interactions, with random phosphoproteomes frequently scoring better and having no distinguishable difference in their ability to identify protein-protein level interactions.

## ACKNOWLEDGEMENTS

Research reported in this publication was supported by the National Institute Of General Medical Sciences of the National Institutes of Health under Award Numbers R35GM138127 and T32GM008715 (to Reagan Portelance). The content is solely the responsibility of the authors and does not necessarily represent the official views of the National Institutes of Health. We would like to thank Dr. Casim Sarkar and Dr. Wesley Errington for sharing the Matlab code and giving guidance on model adaptations. We would additionally like to thank Devon Semoy and Dr. Neel Shah for their helpful guidance in native PAGE for the separation of phosphospecies. The authors acknowledge Research Computing at The University of Virginia for providing computational resources and technical support that have contributed to the results reported within this publication. URL: https://rc.virginia.edu

