## Supplementary Figures for "Structure-informed theoretical modeling defines principles governing avidity in bivalent protein interactions"

### Supplementary Materials for Portelance, Wu, Kandoor, and Naegle

***This PDF file includes:.*** Table S1
Figures S1 to S13
Captions for Data S1 to S4

| Pair | $l_p$ = 4Å | $l_p$ = 60Å | $l_p$ = 70Å | $l_p$ = 80Å |
|---|---|---|---|---|
| 1016/1138 | 20.1812 | 65.1425 | 84.7675 | 110.6075 |
| 1138/1197 | 26.8613 | 57.085 | 72.835 | 94.255 |
| 998/1138 | 28.0388 | 37.715 | 42.225 | 48.365 |
| 1172/1197 | 32.4263 | 34.205 | 34.485 | 34.745 |
| 998/1016 | 33.2287 | 33.785 | 33.815 | 33.845 |
| 1016/1092 | 35.8537 | 223.605 | 329.045 | 489.305 |
| 1138/1172 | 37.5688 | 47.155 | 50.795 | 55.315 |
| 998/1092 | 42.8937 | 341.94 | 501.66 | 732.82 |
| 1016/1172 | 48.1988 | 471.62 | 872.46 | BNR |
| 1092/1172 | 65.3688 | 458.17 | 668.79 | 983.59 |
| 1092/1138 | 121.2613 | 550.685 | 868.525 | 1442.2 |
| 1092/1197 | 134.5437 | BNR | BNR | BNR |

**Table S1. Persistence length effects on predicted $K_{D,B}$.** Pairs are sorted from highest to lowest affinity based on initial model (4Å) predictions. Values are given in nM the predicted affinity of the bivalent species and BNR indicates Bivalency Not Reached in the model. Persistence length ($l_p$ given in angstroms) was initially set to highly flexible (4Å) and 60-80Å represents a range of values fit after experimental values.

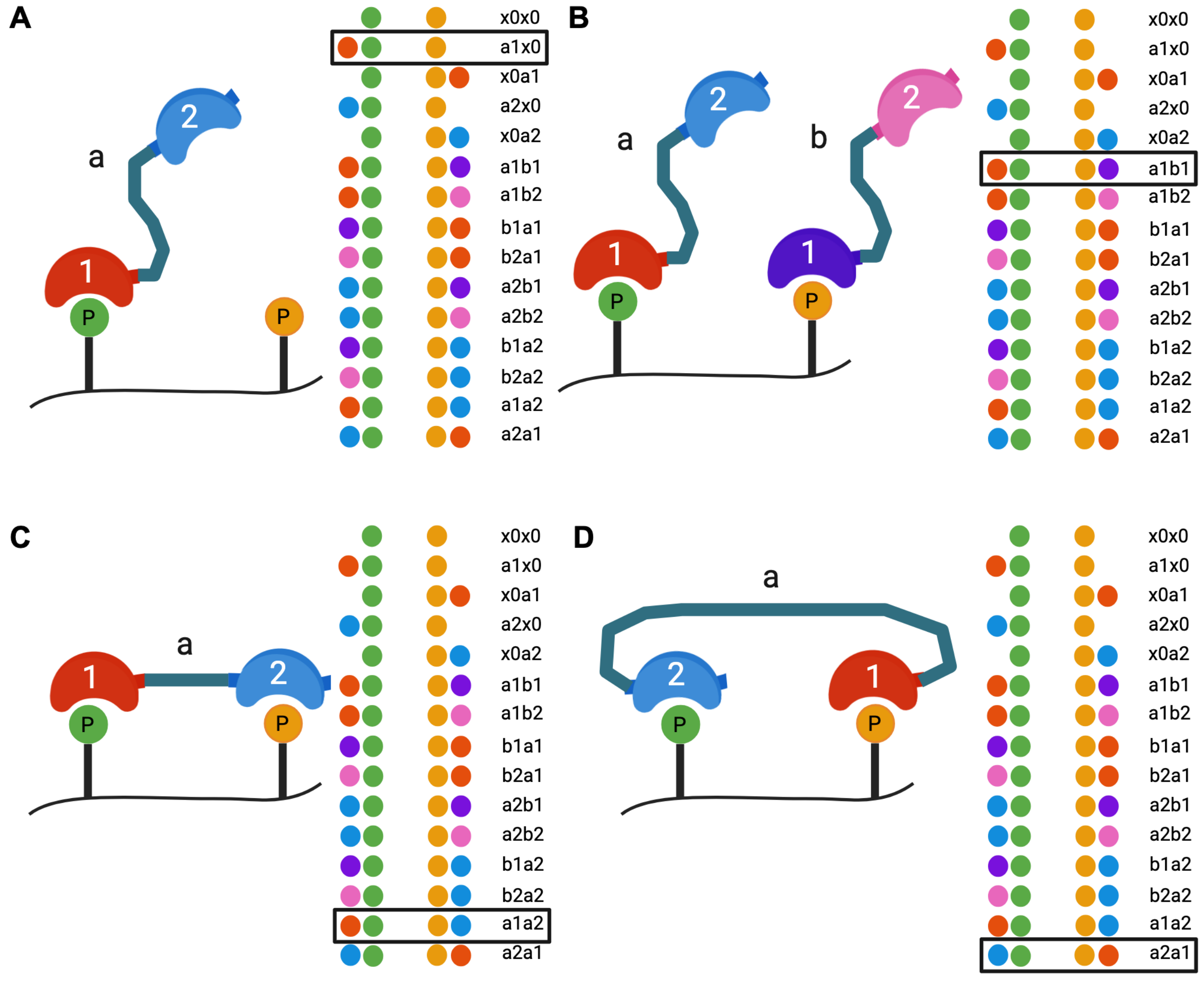


**Fig. S1. Bivalent binding states.** There are 15 possible binding states for a bivalent system. In addition to the unbound state this includes (**A**) four monovalently bound states, (**B**) eight bivalently bound states spanning across two tandem SH2 proteins, and two versions of a bivalently bound state - one (**C**) inline and one (**D**) twisted.

Effect of cross-reactivity with high monovalent or mismatched monovalent affinities

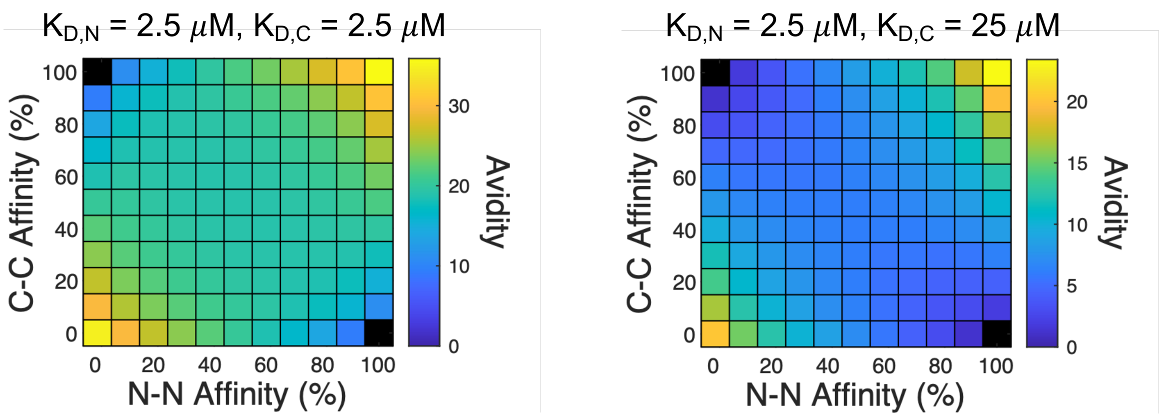


**Fig. S2. Cross-reactivity negatively impacts bivalent interactions.** In order to test for the impact of cross-reactivity we simulated two interactions with fixed monovalent affinity – either as both high affinity (left) or as one high affinity and one low affinity interaction (right). We then changed the cross-binding interaction affinity from N-SH2 to C-pTyr and C-SH2 to N-pTyr, increasing it from no cross-reactivity up to 100% cross-reactivity while decreasing the N-SH2 to N-pTyr and C-SH2 to C-pTyr affinity accordingly to maintain a constant $K_{D,N}$ and $K_{D,C}$.

***Supplementary Figures.***

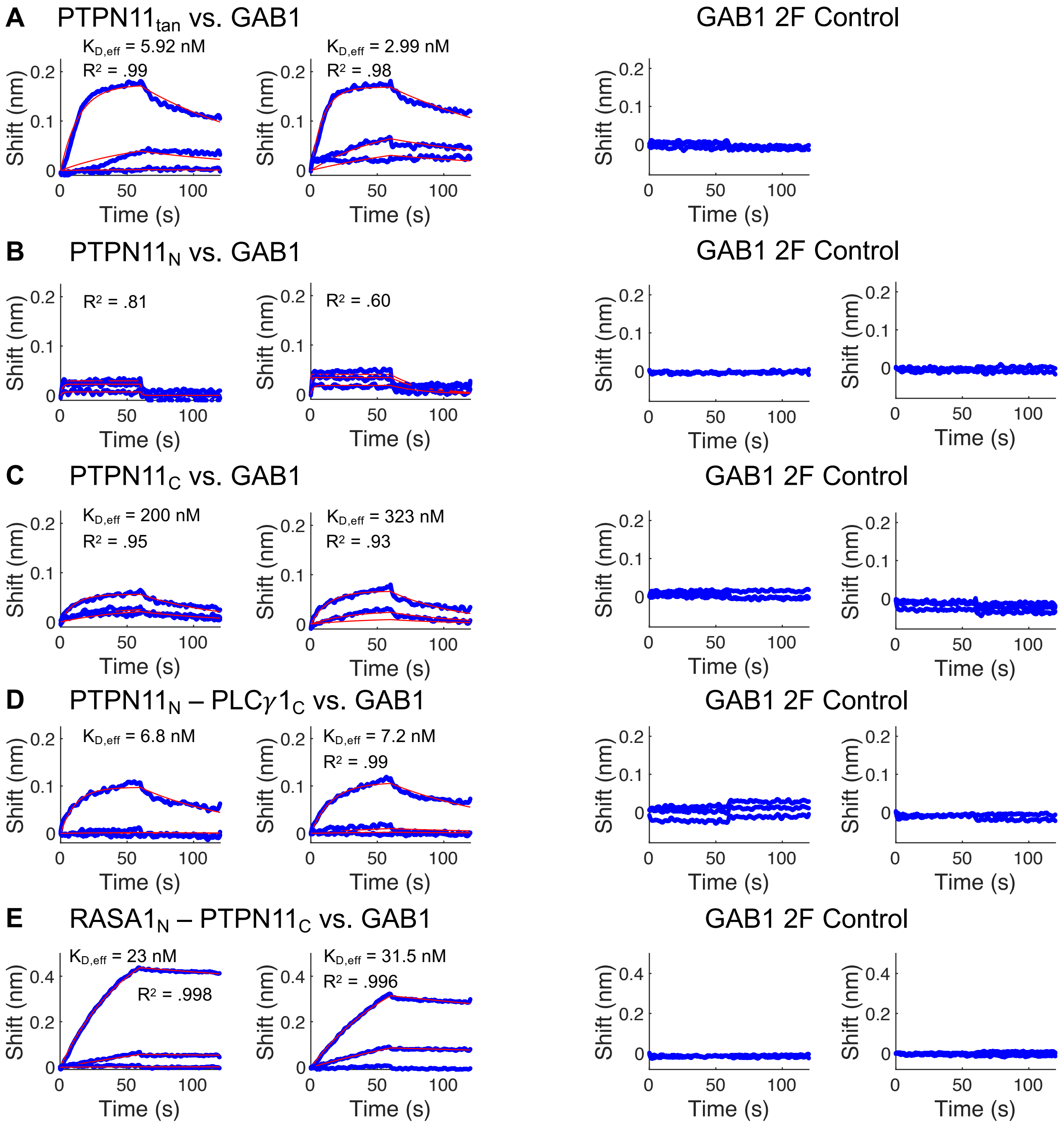


**Fig. S3. Replicates and Y2F controls for BLI experiments of PTPN11 interacting with GAB1.** GAB1 refers to the same preparation of phosphorylated GAB1 C-terminal tail spanning Y627 and Y659. GAB1 2F refers to the negative control with both tyrosines mutated to phenylalanine. The replicate data and the effective $K_D$ are given for (**A**) Tandem SH2 domains of PTPN11 (40, 4, 0.4 nM) (**B**) The N-terminal SH2 domain of PTPN11 (1000, 250, 62.5 nM) (**C**) The C-terminal SH2 domain of PTPN11 (1000, 250, 62.5 nM) (**D**) The chimera of PTPN11 N-terminal and PLC$\gamma$1 C-terminal domain (40, 4, 0.4 nM), and (**E**) The chimera of RASA1 N-terminal domain and the PTPN11 C-terminal domain (400, 40, 4 nM (left) and 200, 50, 12.5 nM (right)).

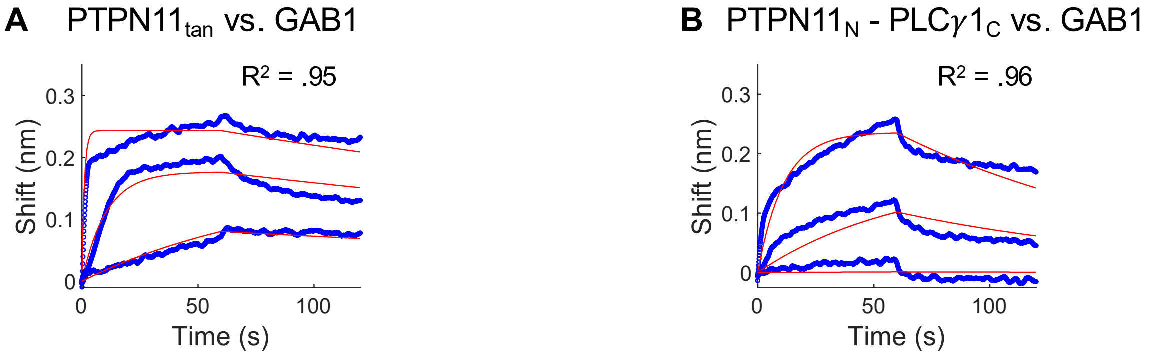


**Fig. S4. Examples of serial dilution range that produces nonlinearities.** Examples of (**A**) PTPN11$_{\mathrm{tan}}$ and (**B**) PTPN11$_{\mathrm{N}}$-PLC$\gamma$1$_{\mathrm{C}}$ vs. GAB1 BLI experiments with too high SH2 concentrations (400, 40, 4 nM) resulting in poor fit.

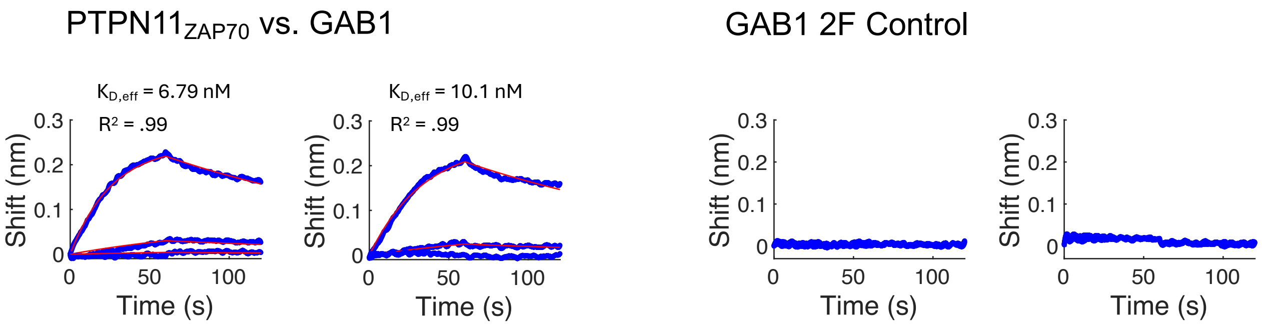


**Fig. S5. Replicates and Y2F controls for BLI experiments of PTPN11$_{\mathrm{ZAP70}}$ binding with GAB1.** GAB1 refers to the same preparation of phosphorylated GAB1 C-terminal tail spanning Y627 and Y659. GAB1 2F refers to the negative control with both tyrosines mutated to phenylalanine. The replicate data and the effective $K_{\mathrm{D}}$ are given for tandem SH2 domains of PTPN11$_{\mathrm{ZAP70}}$ (40, 4, 0.4 nM).

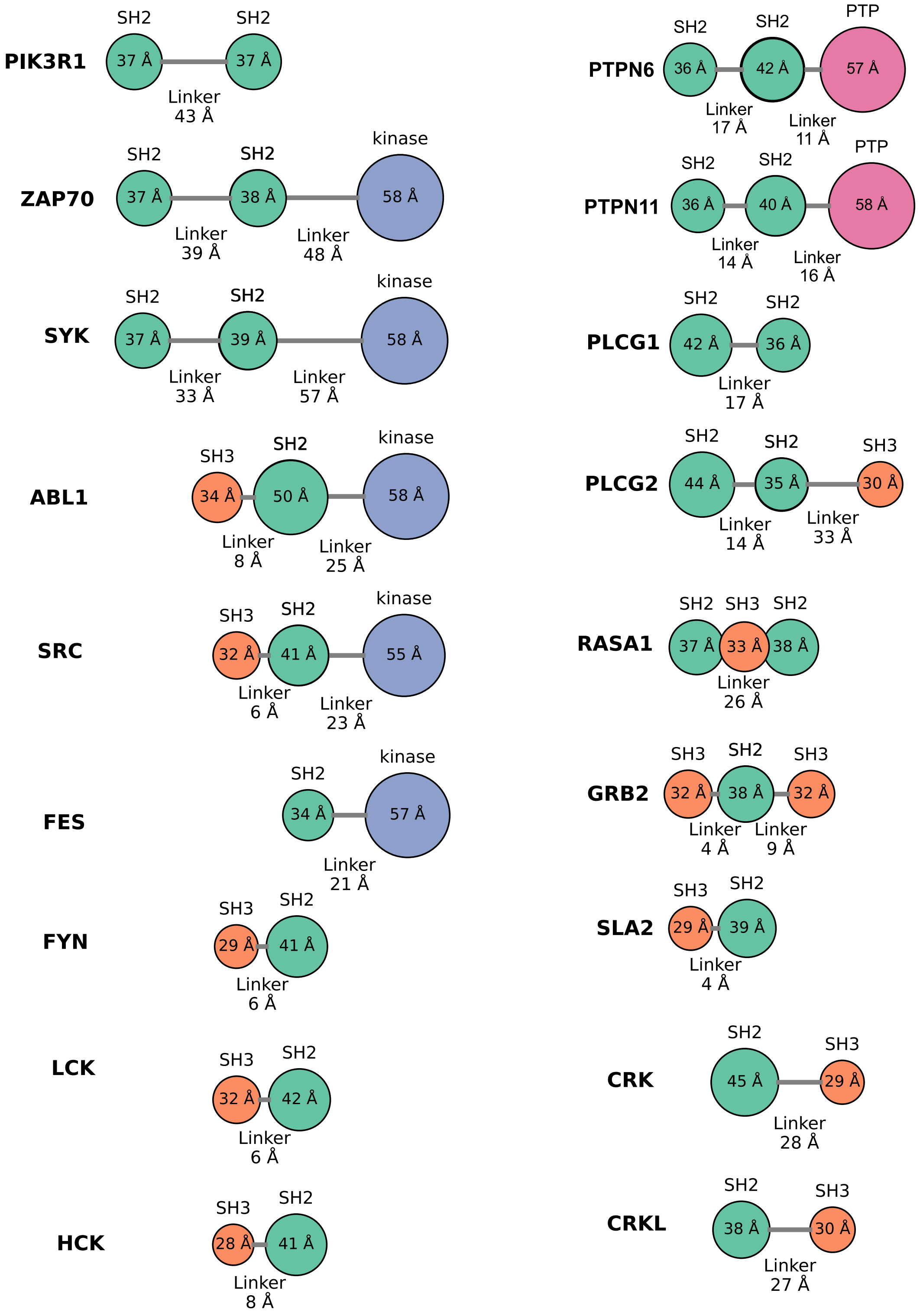


**Fig. S6. Domain architecture of SH2 containing proteins.** The domain architecture of proteins used to measure the linker distance between SH2, SH3, PTP, and Kinase domains is provided along with the size of each domain and distance across each linker. Domains are only included if a linker distance could be measured in an available PDB structure (i.e. both domains are included in a single structure).

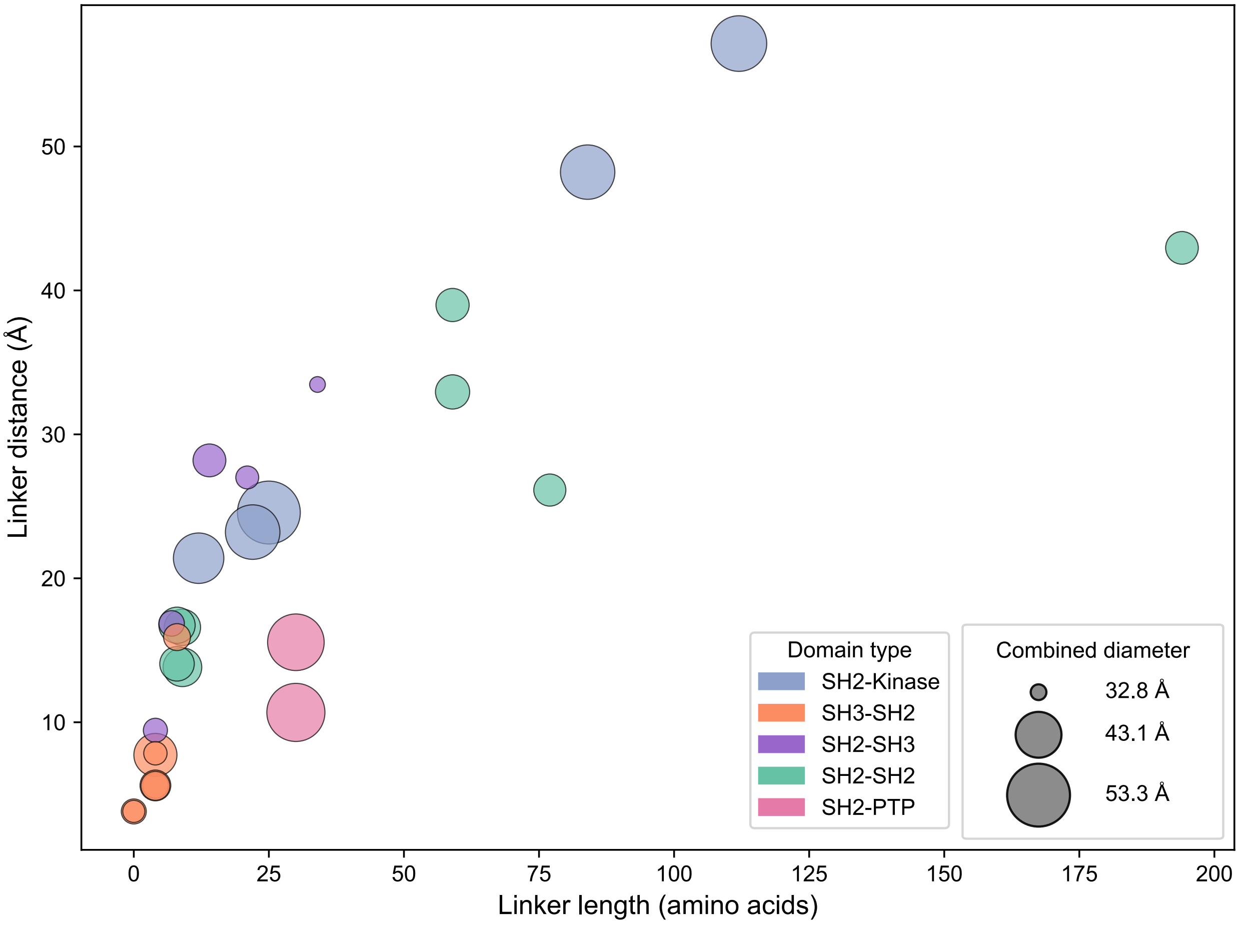


**Fig. S7. Linker size between SH2, SH3, PTP, and Kinase domains.** All linker distances between SH2, SH3, PTP, and Kinase domains in the human proteome were measured if PDB structures were available. The average distance for each protein was plotted against the number of amino acids comprising the linker. The size of the dot represents the possible occlusion size of the linker calculated by summing the radii of each domain bookending the linker.

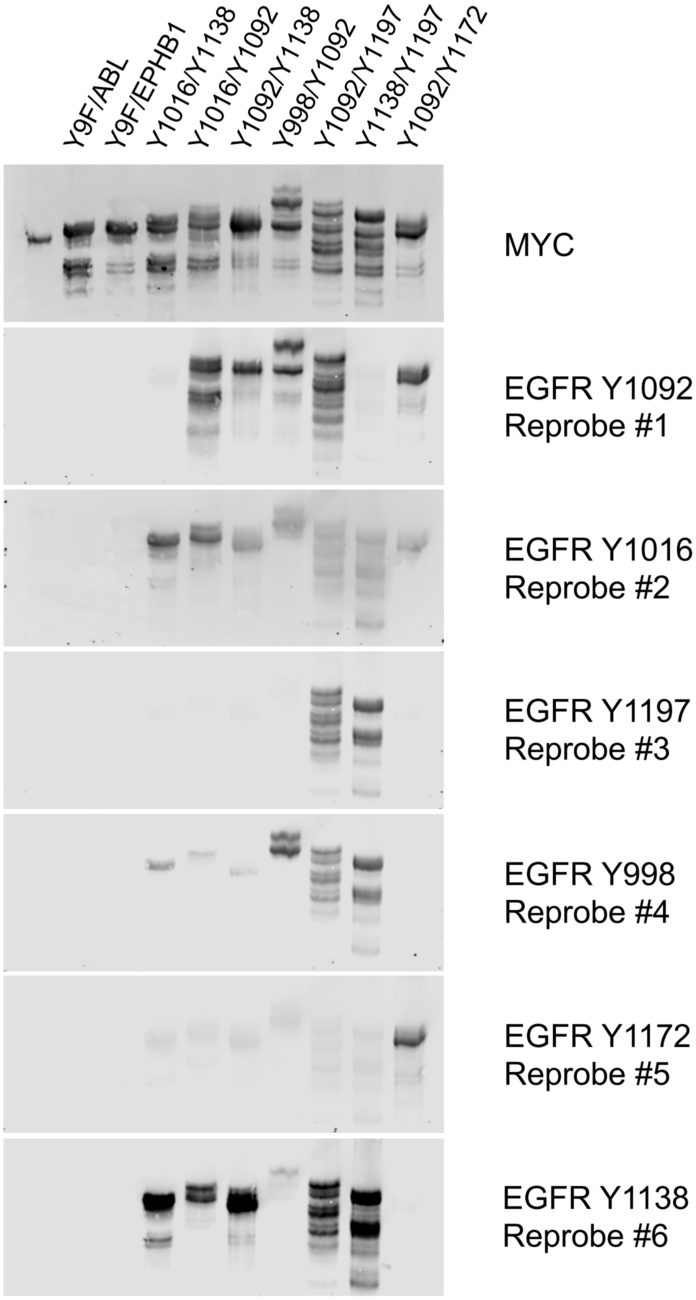


**Fig. S8. Validation of double phosphorylation on EGFR Y7F constructs.** All EGFR Y7F pairs were run on a Zn-Phos-Tag gel and co-probed with anti-MYC antibody and site-specific antibodies for EGFR pTyr sites 998, 1016, 1092, 1138, 1172, and 1197. The anti-ERBB2 Y1196 antibody was used for site 1138 due to homology between the sites, but the antibody proved to be nonspecific and bound broadly across the pairs. The 998 antibody also demonstrated some nonspecificity, and there was some incomplete stripping during the 1016 probe. Clear separation of phosphospecies were seen in some pairs such as Y998/Y1092 while other pairs such as Y1016/Y1138 showed little separation. It is difficult to conclusively determine the extent of double phosphorylation on any individual pair due to variation in migration, but we did show that both EGFR sites exhibited some degree of phosphorylation in each pair. Whether this is a mixture of singly-phosphorylated species or a properly doubly-phosphorylated species remains unclear.

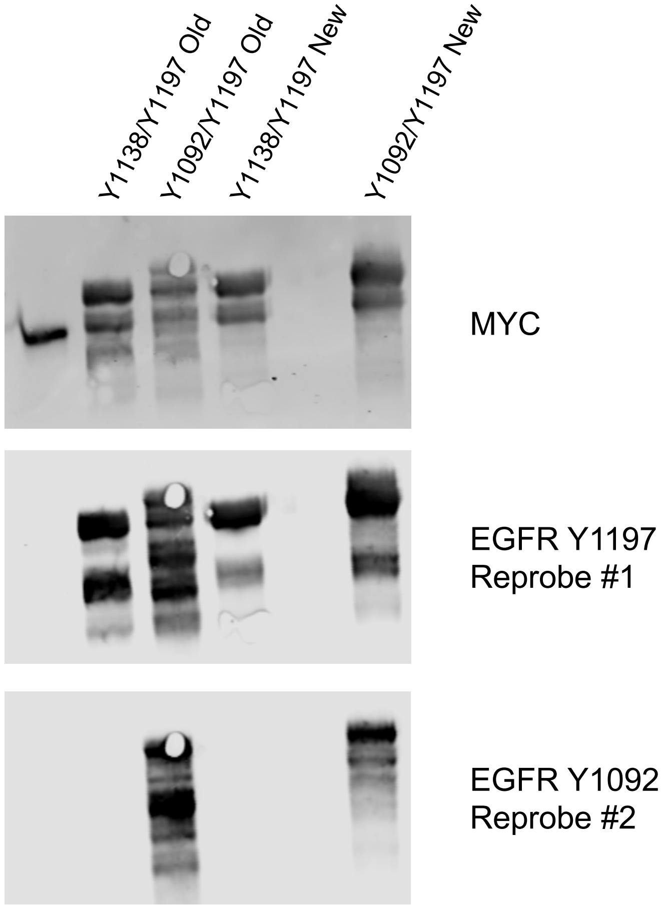


**Fig. S9. Phos-Tag of New EGFR Y7F Constructs.** EGFR Y7F pairs Y1092/Y1197 and Y1138/Y1197 were remade due to the presence of breakdown products in the Phos-Tag gel and run on a new Phos-Tag along with the old version as a comparison. They were probed with anti-MYC antibody and site-specific antibodies for EGFR pTyr sites 1092 and 1197. The anti-ERBB2 Y1196 antibody used for site 1138 was not utilized due to nonspecific binding.

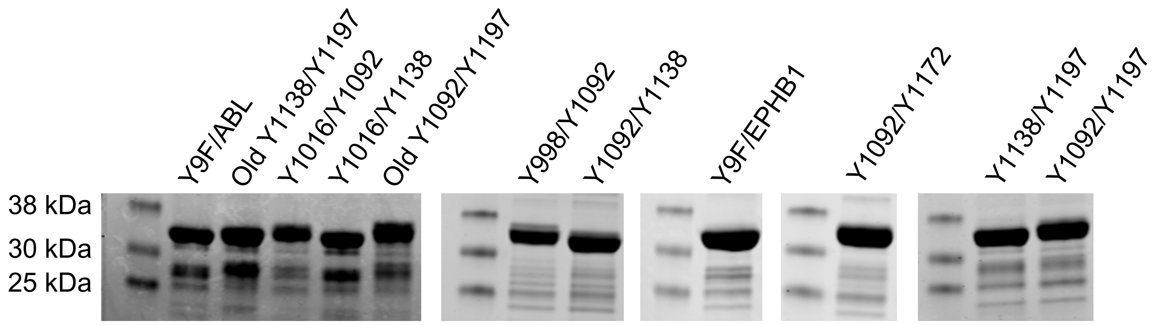


**Fig. S10. Quantification of EGFR Y7F Constructs.** All EGFR Y7F pairs were run on a Coomassie gel to stain for total protein. The final iterations of each pair are provided along with iterations of 1092/1197 and 1138/1197 that were not the final constructs used due to insufficient purity.

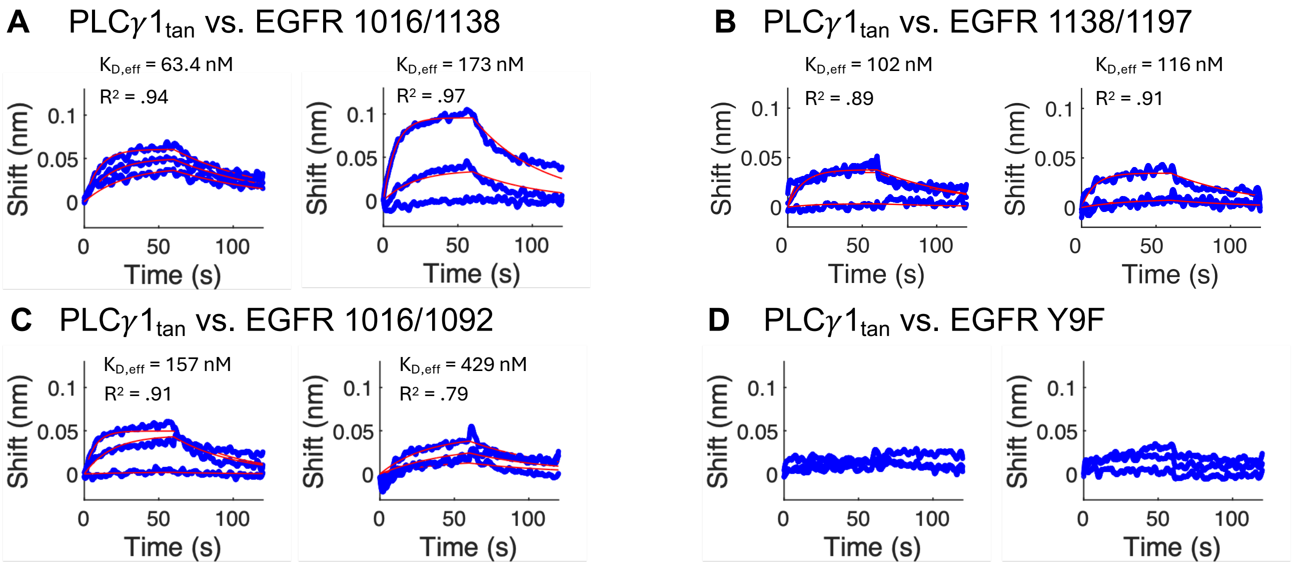


**Fig. S11. Replicates and Y9F controls for EGFR BLI experiments.** EGFR refers to the different preparations of phosphorylated EGFR C-terminal tail with unique pairs of pTyr sites available and the remaining seven pTyr sites mutated to a phenylalanine. EGFR Y9F refers to the negative control with all tyrosines mutated to phenylalanine. The replicate data and the effective $K_D$ are given for (**A**) EGFR 1016/1138 (400, 200, 100 nM (left) and 800, 200, 50 nM (right)), (**B**) EGFR 1138/1197 (625, 250, 100 nM (left) and 500, 200, 80 nM (right)) (**C**) EGFR 1016/1092 (800, 200, 50 nM (left) and 400, 200, 100 nM (right)), and (**D**) EGFR Y9F (800, 200, 50 nM).

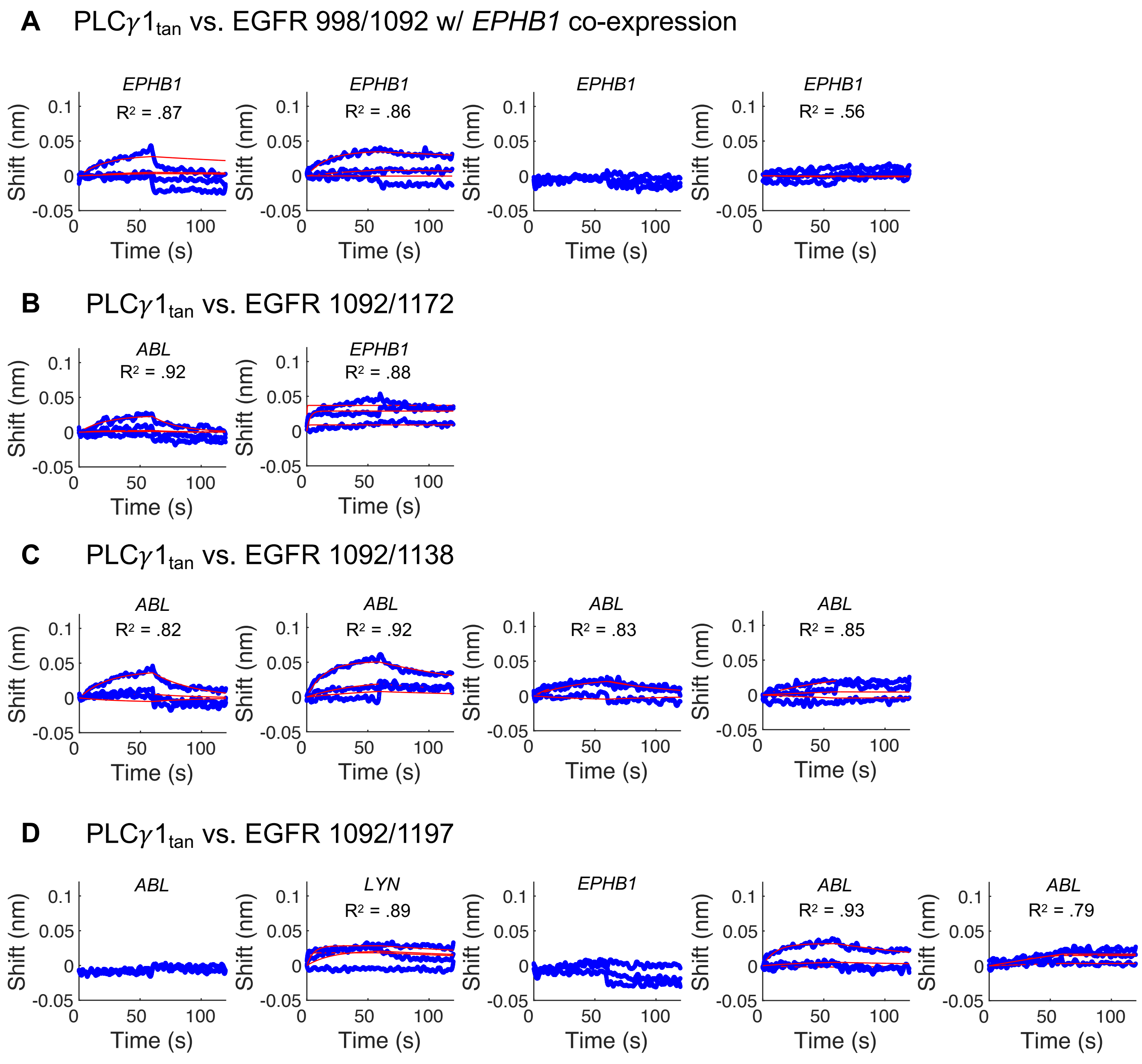


**Fig. S12. EGFR Y7F Pairs with Poor BLI Fit.** In addition to EGFR Y7F pairs 1016/1138, 1138/1197, and 1092/1197, four additional pairs were produced and tested against PLC$\gamma 1_{\mathrm{tan}}$ using BLI. These four pairs - (**A**) 998/1092, (**B**) 1092/1172, (**C**) 1092/1138, and (**D**) 1092/1197 - did not achieve an adequate fit across any replicate tested so a $\mathrm{K}_{\mathrm{D,eff}}$ could not be reported. For certain pairs, multiple kinases co-expressions were tested, and the kinase used for each replicate is listed.

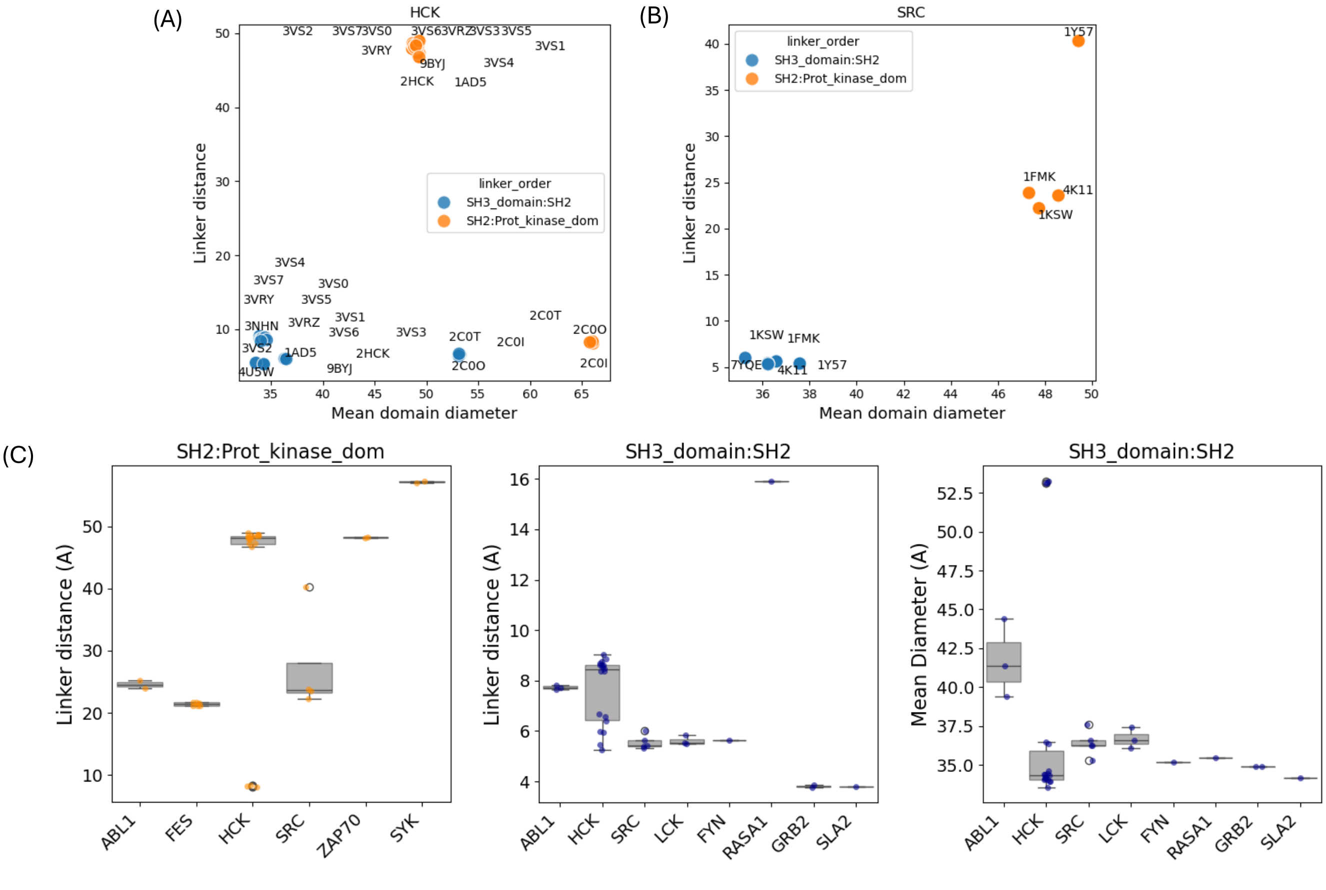


**Fig. S13. Assessing outlier structures within each domain-domain linker group.** Scatter plots showing the mean domain diameter of the two domains (x-axis) versus the linker distance (y-axis) for (**A**) HCK and (**B**) SRC structures. Each datapoint corresponds to an individual PDB structure and is annotated with its PDB identifier. (**C**) Box plots summarizing linker distances and mean domain diameters for domain-domain linker pairs with multiple available structures, grouped by gene.

***Supplementary Data.***

**Supplementary Data S1: Model parameters used for all SH2 domain predictions.** The model parameters used (in parameters.m) and the changes in EffC_Calculator.m used for linker parameters are given for each and every prediction provided in this work.

**Supplementary Data S2: Structure-based extraction of tandem SH2 domain linkers and SH2 domain diameters.** Sheet 1 of this Excel workbook contains all PDB's analyzed for linker distances according to methods, and the average value derived that was used in modeling. Sheet 2 contains the analysis of all SH2 domain diameters in structures used and a final diameter used in the model that is the average of all values.

**Supplementary Data S3: Structure-based extraction of all SH2-relevant linkers.** This workbook contains the extraction of all related PDB files that had an SH2 domain with coverage of a second, or more, domain of interest (phosphatase catalytic domain, kinase catalytic domain, or SH3 domain). PDB_filtered indicates the final structures used, eliminating structures with an inhibitor bound, or other outlier issue as indicated in Methods. Domain_linker_summary sheet is the final average linker distance and domain diameters for all pairs of domains containing an SH2 domain across the PDB. Sizes are given in angstroms, and AA indicates length in number of amino acids.

**Supplementary Data S4: Bivalent pTyr resource.** This workbook contains the full human bivalent pTyr pairs according to methods provided. Sheets labeled Ronan bivalent pTyr and Martyn bivalent pTyr are the reference pairs that had annotated data in the Martyn (44) or Ronan (25) resources. The pair-level data was summarized in Martyn PPI and Ronan PPI sheets – indicating protein-protein interaction level data as accumulated form the pairs of sites.
